# CartoCell, a high-content pipeline for 3D image analysis, unveils cell morphology patterns in epithelia

**DOI:** 10.1101/2023.01.05.522724

**Authors:** Jesús A. Andrés-San Román, Carmen Gordillo-Vázquez, Daniel Franco-Barranco, Laura Morato, Cecilia H. Fernández-Espartero, Gabriel Baonza, Antonio Tagua, Pablo Vicente-Munuera, Ana M. Palacios, María P. Gavilán, Fernando Martín-Belmonte, Valentina Annese, Pedro Gómez-Gálvez, Ignacio Arganda-Carreras, Luis M. Escudero

## Abstract

Decades of research have not yet fully explained the mechanisms of epithelial self-organization and 3D packing. Single-cell analysis of large 3D epithelial libraries is crucial for understanding the assembly and function of whole tissues. Combining 3D epithelial imaging with advanced deep learning segmentation methods is essential for enabling this high-content analysis. We introduce CartoCell, a deep learning-based pipeline that uses small datasets to generate accurate labels for hundreds of whole 3D epithelial cysts. Our method detects the realistic morphology of epithelial cells and their contacts in the 3D structure of the tissue. CartoCell enables the quantification of geometric and packing features at the cellular level. Our Single-cell Cartography approach then maps the distribution of these features on 2D plots and 3D surface maps, revealing cell morphology patterns in epithelial cysts. Additionally, we show that CartoCell can be adapted to other types of epithelial tissues.

**MOTIVATION:** A major bottleneck in developing neural networks for cell segmentation is the need for labor-intensive manual curation in order to develop a training dataset. The present work addresses this limitation by developing an automated image analysis pipeline that utilizes small datasets to generate accurate labels of cells in complex, 3D epithelial contexts. The overall goal is to provide an automatic and feasible method to achieve high-quality epithelial reconstructions and to enable high-content analysis of morphological features, which can improve our understanding of how these tissues self-organize.

## INTRODUCTION

Analysis of epithelial tissue properties at the cellular level has enabled advancement in the understanding of different cellular phenomena during morphogenesis. Traditionally, most approaches were based on two-dimensional (2D) analysis of the apical surfaces of monolayer epithelia. However, the need to understand the 3D morphology of epithelial cells to study organogenesis ^1–4^, cell migration ^5^, branching formation ^6^, tumorigenesis ^7^ or wound healing ^8^ has become evident in recent years. A major breakthrough was the discovery that epithelial cells can present very complex geometries due to the exchange of neighbors along the apico-basal axis. These cell shapes have been called scutoids, and it has been shown that they have a role in morphogenesis as well as in the connectivity and biophysical properties of tissues, cushioning and minimizing cell surface tension and leading to a balanced energetic state ^9–16^. Scutoids represent a new paradigm, but they set a challenge for the quantitative analysis of the complex epithelial 3D packing requiring a very accurate reconstruction of 3D epithelial tissues from microscopy images to allow capturing precise cell shapes and neighboring relationships.

In the last few years, deep learning has become the state-of-the-art solution for the analysis of biomedical images ^17–19^. Deep learning is a subdomain of machine learning that makes use of large (or so-called *deep*) artificial neural networks to solve a wide variety of tasks. As opposed to conventional algorithms, before they can be used, deep learning methods (or models) need to be *trained*. In other words, models can *learn* from a set of examples how to solve a specific task. Once trained, the models can be directly applied to new samples, what is usually called prediction or *inference*. In the particular case of image segmentation, the training dataset is commonly formed by a set of raw images and their corresponding ground-truth annotations or *labels*. This type of learning framework, with both raw and label images available, is known as supervised learning. Furthermore, realistic 3D reconstruction of epithelial cells requires assigning each individual cell a unique label, in a process called *instance segmentation*.

The 3D instance segmentation of microscopy data is a difficult task, especially in the presence of a dense concentration of cells and anisotropic voxel resolution, as it is common in volumetric images of epithelial tissue. State-of-the-art learning-based methods tackle these challenges using a top-down strategy, by first training a deep neural network (DNN) to predict representations of the objects of interest (cells in our case), and then extracting individual instances from those representations using different post-processing methods. Common representations include cell masks or boundaries ^20–23^, distance or flow maps ^24–26^, or a combination of some of the latter ^27,28^. On top of those representations, cell instances are then calculated usually by means of watershed ^29,30^ or graph-partitioning methods ^31–33^. Other techniques have shown success in segmenting cell nuclei and tracking cell lineage ^34,35^, but they do not have the high level of accuracy in cell shape required to obtain detailed geometric and topological information at the cellular level.

Despite the benefits observed from these supervised approaches, their main drawback is the large number of annotated samples needed to establish a training dataset and obtain reliable performance ^36^. Preparing and processing such a large amount of data manually or semi-automatically is usually tedious and time-expensive. This problem arises both from the acquisition time of high-resolution images, as well as from the labeling of raw images performed by experts in the field, which is usually the main bottleneck of the protocol. To address this issue, a common strategy consists in using data augmentation, i.e. synthetically increasing the size of the training data by morphological and intensity transformations or noise addition ^37,38^. However, data augmentation may not be sufficient to realistically recreate the diversity of image data to be processed. A much less exploited alternative to speed up the segmentation protocol would consist in the use of low-resolution images instead, which are acquired and annotated at a considerably faster pace. Certainly, this option would be ideal if the quality of the output segmented cells remains comparable to that obtained with high-resolution images.

In this article, we image, process, and analyze whole Madin-Darby Canine Kidney (MDCK) 3D epithelial cell cultures ^39^. Despite their simplicity, these cysts have previously been used as a suitable model system to study the establishment of cell polarity and cell junctions ^3,40–43^, epithelial morphogenesis and physiology ^44–51^, tumor progression ^52–55^, and for exploring the constraints on epithelial tissue morphogenesis ^56^. MDCK cysts have provided valuable insights to study more complex systems helping to understand the self-organization in organoids, embryoids and the early stages of mammal development ^56–58^.

In our approach, we use a small training dataset of high-resolution images to subsequently produce a large training dataset of low-resolution images that are automatically segmented. Our method follows a top-down pipeline that makes use of a DNN architecture with multiple cell representations and watershed post-processing to initially segment the epithelial cells as 3D instances. These instances are refined by a second post-processing step: a 3D Voronoi algorithm that provides realistic 3D epithelial boundaries where cells are in close contact to each other. The algorithm is based on tiling the space between a set of (Voronoi) seeds by proximity, without leaving any gaps among the generated compartments ^59^. These compartments are called Voronoi cells. Honda and colleagues showed that the Voronoi compartmentalization of a 2D space, after using the cell nuclei as seeds, fitted the pattern of cellular contacts found in epithelial surfaces ^60^. In 3D, this approach has been previously used to simulate the shapes of globular cells ^10^. Here, it is key to increase the quality of our cell segmentation results.

In short, we have developed an accessible and fast tool to investigate the complex organization of epithelial tissues. The production of a large number of samples with an accurate segmentation has opened up a new way of 3D high-content analysis. The representation of the extracted features values in each cell provides maps of the cysts at single-cell resolution. Due to the similarities with the practices of making and using maps, we called our approach Single-cell Cartography, and our high-content segmentation method, CartoCell. The simple observation of these maps reveals the presence of cell morphology patterns where cells are distributed following geometric cues. These patterns illustrate how different the cells within the same cyst really are, and how cells with similar characteristics have the tendency to cluster together in specific zones of the cysts. Importantly, the large number of processed individual cells permits us to quantify the frequency of the patterns and even to find hidden traits of organizational features within the 3D structure of the tissue.

## RESULTS

### CartoCell, a high-throughput pipeline for segmentation of 3D epithelial cysts

The realistic analysis of whole epithelial tissues at the cell level is a critical point to a bottom-up understanding of how tissues self-organize during development. In this work, by means of deep learning and image processing strategies, we have developed CartoCell, an automated pipeline (**Figure 1**) to segment and analyze hundreds of epithelial cysts at different stages (**Table S1**) with minimal human intervention. CartoCell is subdivided into five consecutive phases.

**Figure 1.**
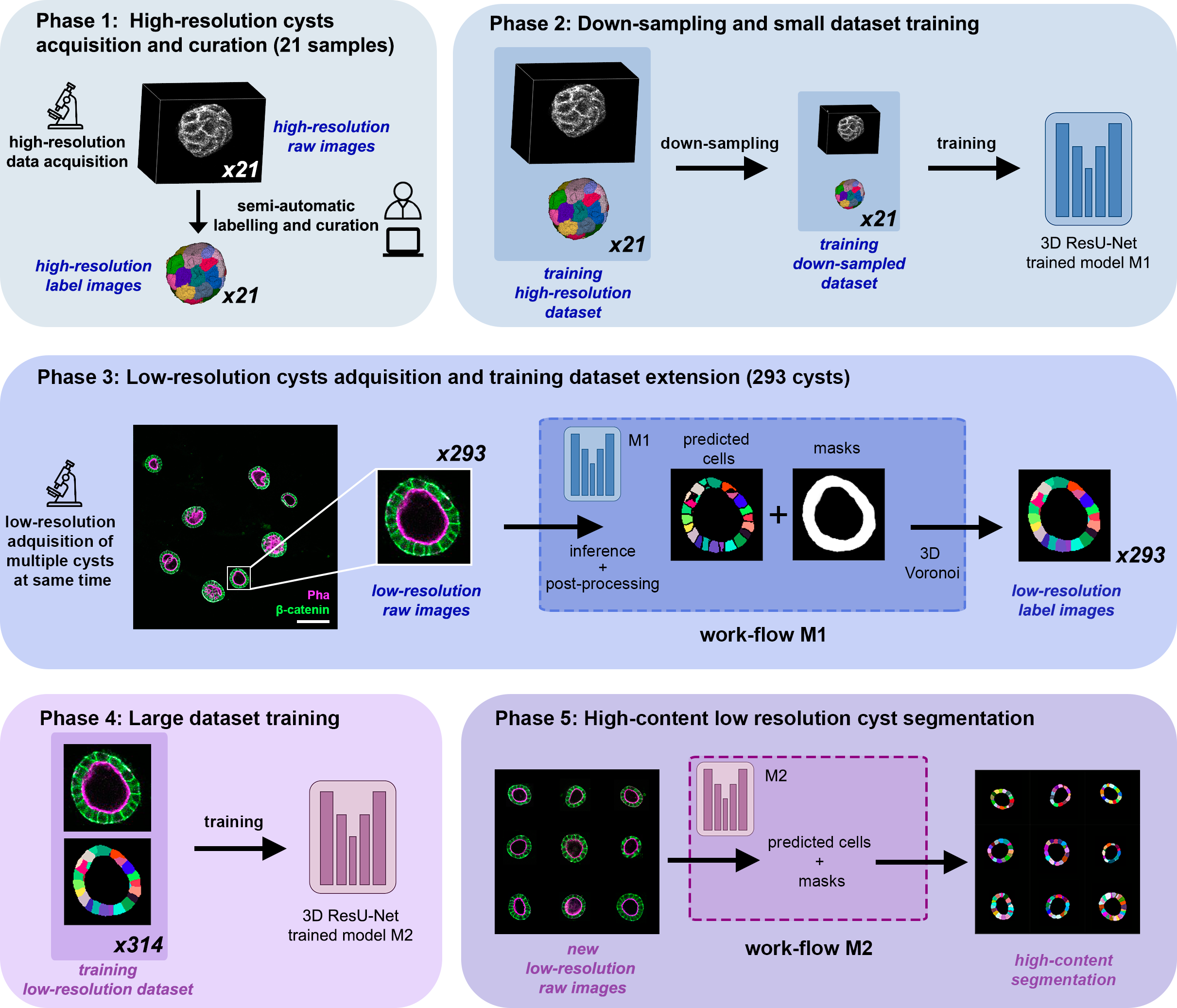
CartoCell pipeline for high-content epithelial cysts segmentation. Phase 1: The “*high-resolution raw images*” consist of confocal Z-stacks images, where the cell membrane is stained. These images are segmented and proofread using LimeSeg, and a custom Matlab code for curation to obtain the “*high-resolution label images*” (**STAR Methods**). Together, the raw and the label images encompass the “*training high-resolution datase*t”. Number of samples = 21. Phase 2: The “*training high-resolution datase*t” is down-sampled to obtain the “*training down-sampled dataset”,* which is the training set for the “model M1”. Phase 3: low-resolution images are obtained from confocal Z-stacks images, stained in a similar way to Phase 1; scale bar, 100 µm. Number of samples = 293. Next, the “work-flow M1” is applied: Inference using “model M1” and subsequent post-processing to obtain individual cell instance predictions and cell masks, followed by the 3D Voronoi algorithm to guarantee that predicted cells remain in close contact. As a result, the “*low-resolution label images”* are generated. Phase 4: Training of the “model M2” on the large “*training low-resolution dataset”*. Number of samples = 314. Phase 5: High-content segmentation of new low-resolution images (unseen by the pipeline) using the “work-flow M2”, which is equivalent to the “work-flow M1” but using the “model M2”. See also **Figure S1**, **Figure S2**, **Figure S3**, **Table S2**, **Table S3** and **Table S4**.

In Phase 1, a small dataset of 21 cysts, stained with cell outlines markers, was acquired at high resolution in a confocal microscope (**Figure 1**, 8 cysts of 4 days and 13 cysts of 7 days, **STAR Methods**). Next, the individual cell instances were segmented using LimeSeg ^61^, **STAR Methods**), a semi-automatic segmentation plugin of Fiji ^62^. The final high-resolution label images were the output of a curation process aided by a custom Matlab code (**STAR Methods**). In particular, we implemented a Matlab graphic user interface to facilitate manual deletion, insertion, fusion, and proper profiling of cell instances and lumen segmentations (**STAR Methods**). The use of high-resolution images simplified and improved the accuracy of manual annotations by providing a clearer visualization of the cysts and their structures. On average, we estimate that the segmentation and curation process took 3 to 5 complete working days, of one person, per cyst. The high-resolution images from Phase 1 provide the accurate and realistic set of data necessary for the following steps (see **Discussion**).

In Phase 2, both high-resolution raw and label images were down-sampled to create our initial training dataset (**Figure 1**). The logic of this step is to take advantage of lower storage requirements, faster acquisition and processing time of low-resolution images. Specifically, the image volumes were reduced to match the resolution of the images acquired in Phase 3 (**STAR Methods**). Using that dataset, a first DNN was trained. The DNN employed was a custom stable 3D residual U-Net (3D ResU-Net) ^63^ (**Figure 1** and **Figure S1**). We will refer to this first model as “model M1” (**Figure 1**, **STAR Methods**).

In Phase 3, a large number of low-resolution stacks of multiple epithelial cysts was acquired (**Figure 1**). This was a key step to allow high-content analysis of samples, since it greatly reduces the acquisition time (**STAR Methods**). Here, we extracted the single-layer and single-lumen cysts by cropping them from the complete stack (**Figure 1**, **STAR Methods**). This way, we obtained a set of 293 low-resolution images, composed of 84 cysts at 4 days, 113 cysts at 7 days and 96 cysts at 10 days (**Figure S2**). Next, we applied our trained model M1 to those images and post-processed their output to produce (i) a prediction of individual cell instances (obtained by marker-controlled watershed), and (ii) a prediction of the mask of the full cellular regions (**Figure 1** and **Figure S1**, **STAR Methods**). At this stage, the output cell instances were generally not touching each other, which is a problem to study cell connectivity in epithelia. Therefore, we applied a 3D Voronoi algorithm to correctly mimic the epithelial packing ^59,60^. More specifically, each prediction of cell instances was used as a Voronoi seed, while the prediction of the mask of the cellular region defined the bounding territory that the cells could occupy (**Figure 1** and **Figure S3**, **STAR Methods**). In previous works, cell nuclei were used as Voronoi seeds, leading to less reliable cell outlines since nuclei may not be exactly located at Voronoi centers ^64^. In our approach, the predicted full cell instance masks are used as seeds, producing more accurate results since only the inter-instance space need to be filled by the algorithm. The output of this phase was a large dataset of low-resolution images and their corresponding accurate labels.

In Phase 4, a new 3D ResU-Net model (“model M2”, from now on) was trained on the *“training low-resolution dataset”*, composed of the newly produced large dataset of low-resolution raw images and its paired label images along with the *“training down-sampled dataset”* (**Figure 1**, **STAR Methods**). This was a crucial step, since the performance of deep learning models is highly dependent on the number of training samples.

In Phase 5, model M2 was applied to new low-resolution cysts and their output was post-processed as in Phase 3, thus achieving high-content segmentation of the desired cysts (**Figure 1** and **Figure S2**).

### Optimization of the method

Once the CartoCell pipeline was defined, we performed an automatic screening of parameters of the 3D ResU-Net to optimize the quality of the prediction of models M1 and M2 (**STAR Methods**). To this aim, we elaborated a test set with 60 new low-resolution cysts (20 cysts at 4 days, 20 at 7 days and 20 at 10 days of development), not used in any of the previous training steps (**STAR Methods**). These cysts were semi-automatically segmented and manually curated to obtain their ground-truth labels (**STAR Methods**). The parameter search for the M1 and M2 models aimed to ensure the highest quality of segmentation, based on the comparison between the prediction of the models and the ground-truth labels of the test set (**Table S2**, **STAR Methods**). In addition, this optimization also demonstrated that M2, a model trained with a large number of low-resolution images outperformed M1, a model trained with a small but perfectly segmented dataset (**Table S3**).

### Performance evaluation and comparison with other methods

To test CartoCell against current alternatives, we compared the performance of our segmentation pipeline with that provided by the state-of-the-art approaches StarDist 3D ^25^, Cellpose ^26^ and PlantSeg ^23^ (**Figure S3**, **STAR Methods**). In all cases, the small down-sampled dataset from Phase 2 was used as the training set and the same 60 new cysts were used as the test set (**STAR methods**). Moreover, for the sake of analyzing method robustness and stability, each method was trained ten times under the same conditions. Segmentation metrics were thus provided on average over those ten repetitions (**Table S3**). In summary, CartoCell compares favorably with state-of-the-art alternatives, especially after retraining our model with the new dataset in Phase 4.

As demonstrated by the segmentation metric values of model M1 and M2 (**Table S3**), one of the keys of the success of CartoCell relies in generating a large yet imperfect training dataset, which greatly enhances the prediction accuracy when training a DNN. To measure this improvement, we quantified the number of perfectly annotated cysts that would be required in a single phase to achieve equivalent results to the whole pipeline of CartoCell. In particular, we trained our ResU-Net with 50, 100, 150, 200 and 250 perfectly annotated cysts (**STAR methods**). As a result, we observed that as much as 90 cysts would be needed to match the performance of CartoCell with only 21 cysts as input (**Figure S3**).

Additionally, we demonstrated that other state-of-the-art DNNs can be used in conjunction with CartoCell. By leveraging on the long imperfect training dataset resulting from Phase 2, these methods can be integrated into CartoCell by replacing the model M2 in Phase 4 (**Figure 1**). In line with the findings in our original CartoCell pipeline, the use of the new M2 models improved the segmentation results with respect to those of the model M1 (**Table S3**).

### Management of results: final curation before biological analysis

As a result of this full process, we got 353 segmented cysts (293 low-resolution from Phase 3 + 60 tests). Next, with the purpose of a detailed quantitative analysis of 3D epithelial packing, we performed a semi-automatic final curation to correct small defects in the segmentation (**Figure S3**, **STAR Methods**). As an outcome, we obtained our final ground-truth dataset, with accurate feature values that we used in the biological analysis (**Figure 2**, **Table S1** and **Table S4**). On average, each imperfect cyst took 12 ± 6 minutes to be curated. Thanks to our ground-truth dataset, we could compare the values of the biological features extracted before and after the final curation step, thus measuring their impact on the results. Nevertheless, the user may opt for skipping this semi-manual step and keep a fully-automatic processing pipeline. In our specific case, we automatically selected for analysis the cysts released after Phase 5 whose epithelial monolayer was completely tiled by cells, i.e., without gaps produced by an under-segmentation. Namely, 307 “closed cysts” out of the 353 segmented cysts (**Figure 1** and **Figure S3**). The mean relative error of all the geometric features extracted from the closed cysts was 5.6 ± 4.1%. In the case of connectivity features, the mean relative error was larger: 13.3 ± 13.9%. (**Figure S3**, **Table S4**, **STAR Methods** and **Discussion**).

**Figure 2.**
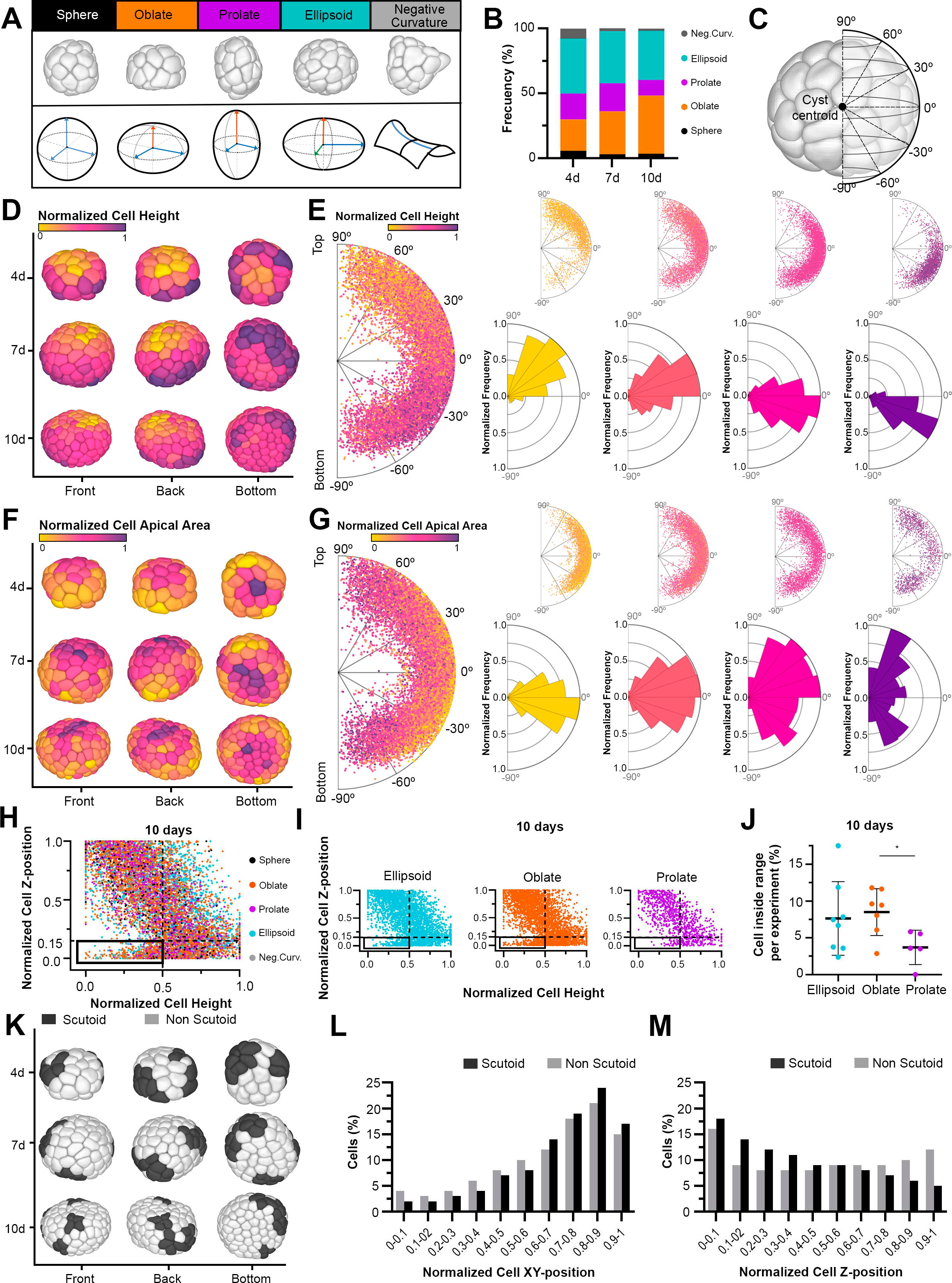
Realistic high-throughput 3D segmentation reveals different morphologies and cell morphology patterns in MDCK cysts. **A**) Shape classification of single-layer, single-lumen cysts. (Top) The 3D rendering of representative segmented cysts. (Bottom) Schematic representation of the morphological cyst classification. **B**) Frequency distribution of the different cyst shapes at 4, 7 and 10 days. **C**) Schematic representation of the relation between 3D reconstructions of the cyst and the 2D plots. The drawing shows how the position and angle of the cell centroids is represented. **D-E**) Single-cell Cartography of “cell height” feature. **D**) Computer rendering of three representative segmented cysts: front, back and bottom views. The cell color scale symbolizes the value of the cell height (normalized by cyst). **E**) (Left) Polar scatter showing the normalized distance and angle of cell centroids regarding the cyst centroid (considering 0 degrees any vector contained in the XY plane of the cyst centroid; positive angles correlate with cell centroids placed over the cyst centroid, and cells located below the cyst centroid XY plane are represented with negative angles, as indicated in panel **C**), with a heatmap coloring the normalized value of cell height. 20391 cells from 353 cysts were represented. (Right) Cell sorting by the normalized cell height values, from left to right: 0-0.25, 0.25-0.5, 0.5-0.75. 0.75-1. On top, a scatter polar diagram showing the angle and distance of cell centroids regarding the cyst centroid, with a heat map coloring based on the quantification of the normalized cell height; on bottom, a polar histogram accounting for the frequency of cell positions. **F**, **G**) Single-cell Cartography of the “apical area” feature. The same color code and plotting properties as panels **D**-**E**. **H-J**) The cyst morphology affects the cartography of cell height at 10 days. **H**) Cell Z-position versus cell height (normalized per cyst). The color of the dots is determined by the kind of cyst the cells belong to. 8528 cells from 116 cysts of 10 days were analyzed. **I**) Scatter plots showing the comparison in cell Z-position versus cell height in 10-day ellipsoid, oblate and prolate cysts. **J**) Cell proportion in the highlighted quadrant of normalized cell Z-position < 0.15 and normalized cell height <0.5 in 10-day ellipsoid, oblate and prolate cysts. The percentage is computed with respect to the total number of cells for each morphology and experiment. Data are represented as mean ± SD. * p value <0.05 (t-Student test). **K-M**) Single-cell Cartography of scutoids. **K**) Computer rendering of three representative segmented cysts in 3 perspectives, showing scutoids in black and non-scutoidal cells in light gray. **L**) Histogram representing the cell proportion at different intervals of the normalized XY-position distance with respect to the XY coordinate of the cyst centroid. Percentage of all cells from the 353 segmented cysts. **M**) Histogram representing the cell proportion of scutoidal and non-scutoidal cells at different intervals of the Z-position distance with respect to the Z coordinate of the cyst centroid. Percentage of all cells from the 353 segmented cysts. See also **Figure S4**, **Figure S5**, **Figure S6**, **Table S1** and **Table S5**.

### Epithelial cysts adopt different shapes in 3D culture

The pipeline that we have developed is able to realistically reconstruct the whole cyst and its lumen. In this way, it allowed us to identify the real shape of the complete 3D epithelial structure. We found that the full set of processed cysts (our “*ground-truth dataset*”) can present a high heterogeneity in terms of shape (**Figure S2**). We analyzed the 3D structure of the total number of single-lumen cysts pointing to two geometrical features: axes lengths and solidity (a curvature index, **Figure S4**, **STAR Methods**). We considered two axes to be similar when the difference in their lengths was inferior to 10%. According to these considerations, we established five types of shapes (**Figure 2A** and **Figure S2**, **STAR Methods**). When the three axes of symmetry were similar in length, cysts were classified as *spheres* (4.0%). When 2 of the 3 axes were similar, but one differed, they were called *spheroids*. Moreover, spheroids were divided into *prolate* (18,1%) when the different axis was the major one, and *oblate* when it was the minor one (34.3%). Cysts with three axes of different lengths were classified as *ellipsoids* (39.9%). Finally, we categorized cysts as presenting *negative curvature* (3.7%) when solidity was less than 0.9, independently of axes values (**Figure 2A** and **Figure S4**, **STAR Methods**). We also quantified the frequency of each type of shape at 4, 7 and 10-day cysts. We found that at 4 and 7 timepoints the most frequent shapes were ellipsoids with the percentage of oblate cysts increasing with the time of culture (**Figure 2B**). At 10 days there was a 37.9% of ellipsoids and a 44.8% of oblate cysts.

### Single cell geometric analysis reveals cell morphology patterns in the MDCK cysts

Reconstructing the 3D outlines of all the cells allowed the precise quantification of a large number of geometrical and connectivity characteristics (**Table S1** and **Figure S4**). We designed a strategy to visualize these data in two ways: 3D maps of the surface of the reconstructed cysts and 2D plots that represent the position of all the cells analyzed within the cysts (**Figure 2C**). We named this approach Single-cell Cartography. In the case of the 3D maps, we obtained a readout of seven cell geometric features by plotting their normalized values using a color palette (**Figure 2D, 2F** and **Figure S4**, **STAR Methods**). Regarding the “cell height” there was a clear gradient “top to bottom” with the cells with lower values on the top of the cyst, and a progressive increase of their height towards the base (**Figure 2D**). Importantly, detailed examination of the entire surface of the cysts revealed a subpattern: just in the center of the bottom region, the cells were shorter (**Figure 2D**). These complex cell morphology patterns were consistent among a high number of cysts and appeared at different time points (**Figure 2D**). To confirm that the pattern was general, we leveraged high-content analysis and visualized the height of each cell of all processed cysts. We plotted the “cell height” data (with the color code used in the 3D cysts) on a scatter-polar diagram considering the angle and radius of the cell centroids with respect to the centroid of the whole cyst (**Figure 2E**). In this way, we were able to visualize the distribution of the “cell height” feature in all cells, from all cysts, at the same time. The plots confirmed the patterns observed in the individual cysts: “shorter” cells (yellower colors) were located at the top and center of the bottom of the cysts, while “taller” cells (pink-purple colors) were found at the periphery of the lateral and bottom part of the cysts. A similar cell morphology pattern, although not that evident, was observed in the distribution of “cell basal area”, “cell volume”, and “cell surface area” values (**Figure S5**). Furthermore, we found a different pattern involving the distribution of the “cell apical area” values (**Figure 2F, 2G**). In this case, cells with a bigger apical area were enriched on the top and on the bottom of the cysts. Meanwhile, cells with the smaller apical area were located in the middle region of the cysts. We also found that “cell solidity” or “cell aspect ratio” characteristics did not show any pattern (**Figure S5**).

### The cell morphology pattern of some features can correlate with the shape of the whole cysts

Our high-content approach revealed that the MDCK cysts can present different shapes and also intrinsic cell patterns. To test if the cell morphology patterns can be affected in some way by the global shape of the cysts, we plotted the values of the features against the position of the centroid of the cells in the Z-axis of the cyst. In this way, we can quantify differences in populations of cells between the three more abundant categories: ellipsoids, spheroids oblate and spheroids prolate. In the case of “cell height”, there was a clear and robust gradient from “shorter” to “taller” cells, from the top towards the bottom on the three types of shapes analyzed (**Figure 2H, 2I**). However, a more detailed analysis of the bottom-left side of the graphs (corresponding to the shorter cells in the base of the cysts) revealed significant differences on the 10-day cysts (**Figure 2I, 2J**) but not on 4-day and 7-day cysts (**Figure S6** and **Table S5**, **STAR Methods**). We also obtained differences in the case of the “basal area” feature, but again, only on 10-day cysts (**Figure S6** and **Table S5**). Conversely, we did not find differences at any time point with the “cell apical area” feature (**Figure S6** and **Table S5**). Our results suggest that the shape of the whole cyst could correlate with changes in cell morphology patterns (see **Discussion**).

### The emergence of cell packing patterns in the cysts

Motivated by the finding of cell morphology patterns in the distribution of the values of cell geometric features, we examined the presence of particular arrangements linked to the connectivity of the cells. To this aim, we obtained Single-cell Cartography representations of the distribution of scutoids ^11^ in the cysts (**Figure 2K-2M**). In this case, the high heterogeneity in the number of scutoids per cyst and their distribution did not enable the identification of any clear pattern using the 3D reconstructions of the cysts (**Figure 2K**). Then, we plot the total number of cells and analyze the distribution of their position along the XY-axes (**Figure 2L**) and the Z-axis (**Figure 2M**) of the cysts. We did not find differences in the distributions in the first case. However, our analysis detected a significant increase of the proportion of scutoids from top to bottom of the cysts (**Figure 2M**) that was not observed in non-scutoidal cells (Chi-Square test, **Table S5**, **STAR Methods**). Our results suggest that cells pack following self-organization patterns in the MDCK cysts.

### Environmental perturbations can alter cell morphology patterns

After finding cell morphology patterns in MDCK cysts, we wondered if they remained consistent when the cell culture environment was modified. Specifically, we investigated the impact of hypoxia ([O_2_] = 1%) on the architecture and organization of 3D cysts (**Figure 3A**, **STAR Methods**) since it has been shown that hypoxia can affect cystogenesis by altering polarity in MDCK cell cultures ^65^ and even inducing epithelial branching ^66^. In our cultures, we found a significant reduction in cell number and the cyst and lumen sizes throughout the whole experimental observation period (**Figure 3B** and **Table S6**). Then, we employed the Single-Cell Cartography approach to analyze 7729 cells from 206 segmented and curated hypoxic cysts and compared the patterns in both normoxia and hypoxia. We observed similar patterns in the case of “cell height” (compare **Figure 2D, 2E**, **Figure S6** and **Table S6**, **STAR Methods**), as well as “cell basal”, “cell volume”, “cell solidity” and “cell aspect ratio (**Table S6**, **STAR Methods**). However, we found a different pattern on the distribution of the “cell apical area” values (compare **Figure 2F, 2G** and **Figure 3C, 3D**). Under hypoxic conditions, cells with the bigger apical area were enriched at the top and bottom regions (like in normoxia), but also in the middle region of the cysts (**Figure 3D** and **Table S6**, **STAR Methods**). Furthermore, in contrast to normoxia, where significant differences in the distribution along the Z-axis of the cyst were found between scutoid and non-scutoidal cells (**Figure 2M**), distinctions were not observed under hypoxic conditions (**Figure S6** and **Table S6**, **STAR Methods**).

**Figure 3.**
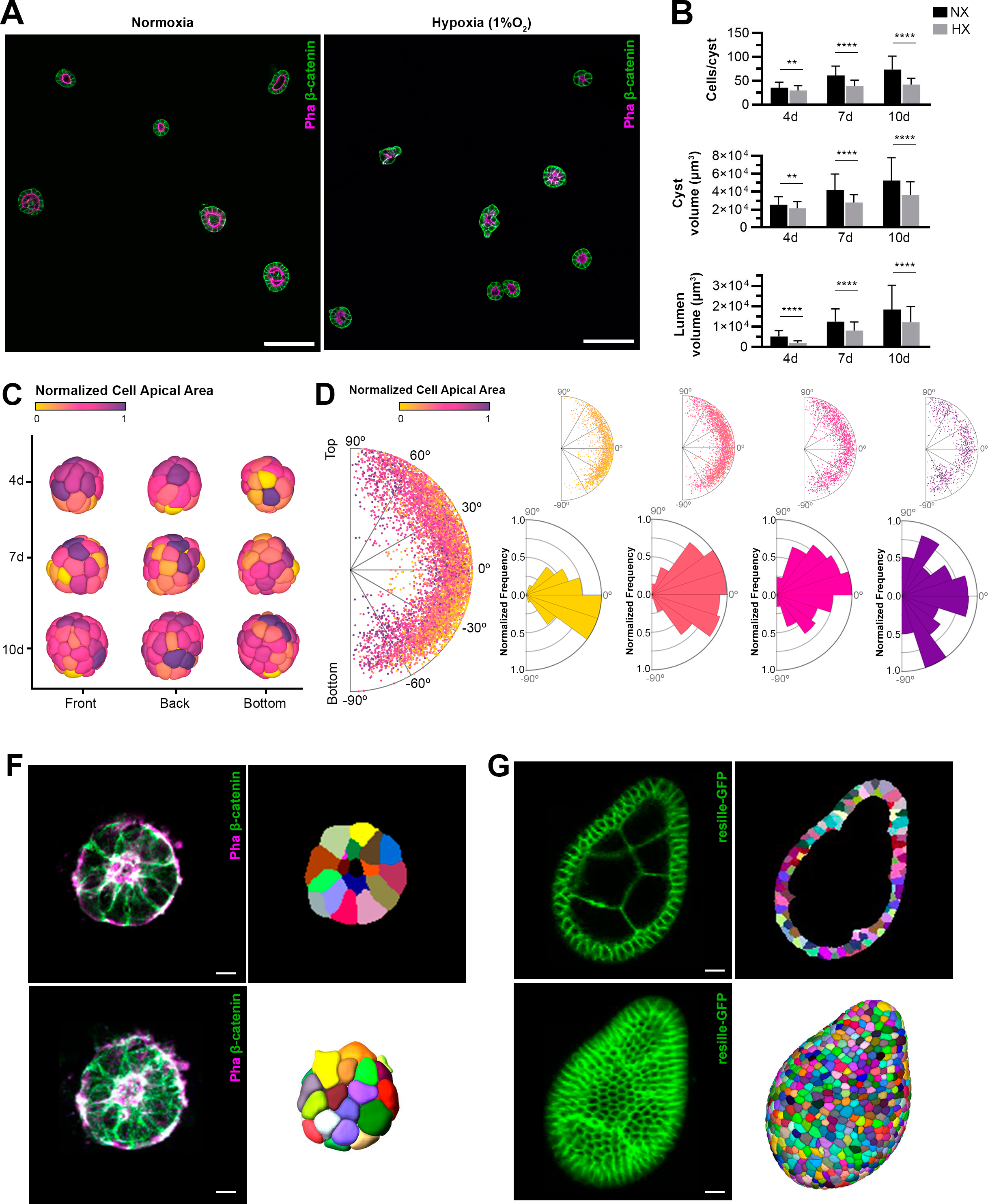
CartoCell segmentation on other epithelial tissue datasets. **A-D**) Hypoxia can induce cell morphology patterns changes in MDCK cysts. **A**) Middle sections of top-to-bottom confocal microscopy Z-stack images of representative cysts cultured under normoxic (left) or hypoxic (1% O_2_) (right) conditions at 4 days. Cell contours were stained with Alexa Fluor 647 phalloidin (magenta) and anti-β-catenin antibody (green) (**STAR Methods**). Scale bars, 100 μm. B) Quantification of cyst size in normoxic (NX, black) and hypoxic (1% O_2_) (HX, grey) cysts at 4, 7 and 10 days. Mean and SD are shown for: number of cells per cyst, cyst volume and lumen volume. Data were obtained from over 60 segmented cysts at each time point, from at least 3 independent experiments. “**” and “****” indicating p ≤ 0.05 and p ≤ 0.001 respectively (Mann-Whitney U test). **C**, **D**) Same representations as Figure 2D**-2G** to map the feature “cell apical area” in hypoxic (1% O_2_) MDCK cysts. 7729 cells from 206 segmented cysts were represented. **F**, **G**) CartoCell segmentation of mouse embryoids (**F**) and *Drosophila* egg chambers (**G**). (Top, left) Middle section of confocal microscopy z-stack images, where in **F**, cell contours were stained with Alexa Fluor 647 phalloidin (magenta) and anti-β-catenin antibody (green) and in **G** with resille-GFP (**STAR Methods**). (Top, right) 2D segmentation of the previous section. (Bottom, left) Half projection of z-stack images with the same stains for cell contours as on top. (Bottom, right) 3D computer rendering. Scale bars, 10μm. See also **Figure S6**, **Table S6** and **Table S7.**

### Applicability of CartoCell to other epithelial models

To assess the adaptability of CartoCell to other epithelial models (differing in morphology and in staining protocols) we evaluated its performance on volumetric confocal images of two different tissues. First, we applied the complete CartoCell pipeline to a dataset of 323 mouse embryoids (**Figure 3F**), which are *in vitro* models derived from embryonic stem cells (ESCs) ^58,67^ (**STAR Methods**). Compared with the MDCK cysts, these embryoids exhibit great variation in terms of cell shape and organization such as a smaller lumen, apoptotic cells, and numerous reverse blebs ^68^. For these reasons, and despite using the same membrane markers as for MDCK cysts, new ground-truth data was generated to feed CartoCell (**STAR methods**). In this scenario, we applied our pre-trained model M2 to generate the initial labels for 21 embryoids. The curation of these embryoids (average curation time = 71 ± 28 min/embryoid) was performed to generate the new training dataset. This approach allows us to achieve rapid and high-quality labeling without the requirement for high-resolution images, resulting in the omission of the downsampling step in the CartoCell pipeline (Phase 2, **Figure 1**). These meticulously curated embryoids constituted the training dataset for model M1. Subsequently, model M1 was trained and used to predict the labels of 282 new embryoids (Phase 3, **Figure 1**). As a result, a substantial number of embryoids with labels that exhibit certain imperfections were obtained. These images, along with the initial 21 embryoids, facilitated the training dataset to train the model M2 (Phase 4, **Figure 1**). This essential balance between a large dataset size and the presence of labels that, while not perfect, remain reliable, culminated in the development of a robust M2 model. To test the model M2, 20 unseen embryoids were inferred and semi-automatically curated using custom software (**STAR Methods**) (average curation time = 61 ± 21 min/embryoid). The comparison between the inference and the ground truth (**STAR Methods**) is provided in **Table S3**, highlighting the precision of CartoCell and illustrating its ability to handle the intricacies of this tissue in comparison to cysts, while maintaining a comparable level of accuracy (**Table S3**).

Second, we tested CartoCell on the well-established *in vivo* model of the *Drosophila* follicular epithelium, which has been widely used to study tissue organization ^69,70^. The follicular epithelium is a cellular monolayer that surrounds developing oocytes forming egg chambers connected to each other to form the ovarioles located in the ovaries ^71^. Instead of using different markers as in cyst experiments, we utilized only one cell membrane marker (Resille-GFP, **Figure 3G**, **STAR Methods**) for visualizing the follicular epithelium of the egg chambers at various stages of development, ranging from early stage 2 to late stage 7. Despite the differences from the original CartoCell set up, we did not require a new training dataset with the egg chamber images to infer the cell prediction from the M2 model (**STAR methods**). We also quantitatively evaluated the resulting segmentation compared to a curated test dataset of 20 egg chambers. The average correction time varies from 45 ± 42 min/egg chamber in early stages to 129 ± 74 min/egg chamber in late stages (**Table S7**, **STAR Methods**). These differences in the curation time were due to the increase in the cell number along the egg chamber development, from 107.6 ± 31.5 cells at early to 853.8 ± 126.2 cells at late stages (**Table S7**, **STAR Methods**). The results obtained showed a high level of accuracy (**Table S3**). These findings provide strong support for the robustness and validity of our model, confirming its capability to effectively address the challenges associated with different tissue types and staining techniques.

## DISCUSSION

In this work, we present CartoCell, a high-content segmentation framework that, coupled with our Single-cell Cartography approach, provides new solutions on the study of 3D complex epithelia. The combination of both tools allows the analysis of hundreds of whole epithelial cysts at the cellular level. We depict the values of any morphological or connectivity parameter at cellular resolution in two ways: using heatmaps of the feature values over 3D reconstructions of each cyst, and with 2D plots of the feature values and the spatial distribution of all the analyzed cells (>20,000 cells).

The generation of a large and sufficiently general training dataset of 3D segmented epithelia is the bottleneck of high-content analysis of epithelial 3D packing. The repositories providing segmented 3D epithelia that could be used as a training dataset are very few and case-specific ^23,72^. For that reason, CartoCell starts with the accurate annotation of a small number of high-resolution samples (Phase 1, 21 cysts, **Figure 1**). Despite being time-consuming, this step facilitates the interactive curation process. As observed by annotation experts, when conducting annotations from scratch, it becomes much easier to identify the actual shape of individual cells (i.e., cell outlines) when working with high-resolution images, contributing to improved annotation quality. Only later, the reduction of resolution is possible while maintaining the precision in the identification of cell outlines (Phases 2 and 3, **Figure 1**). Importantly, the downsampling resolution must match that of the low-resolution Z-stacks acquired in batches of multiple cysts simultaneously. Here, we leverage our deep learning approach to carry out two main steps. First, the small down-sampled dataset is used to train our custom DNN, and subsequently infer a segmentation over hundreds of cysts. Furthermore, thanks to the 3D Voronoi post-processing (**Figure 1** and **Figure S3**), the largest proportion of the segmented cells and their outlines are realistically predicted (**Table S3**). Although these cysts present some little imperfections in their segmentation, they are fundamental for the second step, consisting of retraining our DNN using hundreds of the previously imperfect segmented cysts (also called weak labels ^73^, Phase 4, **Figure 1**). The use of this high number of cysts, despite not being perfectly segmented, adds generality to the model, providing highly reliable results (**Table S3**) even on completely newly acquired cysts (Phase 5, **Figure 1**).

Regarding the usability of CartoCell by the community, here we provide an open-source, well-documented and easy-to-use (without programming skills) segmentation tool that can be used for any lab immediately (see **Data and code availability** section for details and tutorials of CartoCell and Single-cell Cartography). In addition, all the DNN segmentation models generated in this work have been made publicly available. CartoCell outperforms the state-of-the-art alternatives when our DNN model is trained on a small low-resolution dataset (Phase 2, **Figure 1**). The results are substantially better when it is retrained on the larger low-resolution dataset produced by our own pipeline (Phases 3-5, **Figure 1** and **Table S3**). In fact, it is possible to integrate other state-of-the-art segmentation methods with CartoCell, leveraging the benefits of training dataset augmentation by generating realistic but imperfect labels (Phases 1-3, **Figure 1**), and employing other approaches only for the final training and prediction (Phases 4-5, **Figure 1**), with a substantial enhancement in their segmentation quality as we have shown (**Table S3**).

We find that the values of the features extracted from the fully automated segmentation are very reliable when compared with the ground-truth segmentations: only a mean relative error of 5.6% for the geometric parameters (**Figure S3** and **Table S4**, **STAR Methods**). However, the differences were larger in the case of connectivity characteristics, suggesting that the final curation step is necessary for these types of features. Nevertheless, the accurate detection of the epithelial packing and connectivity of the tissues is an increasingly complex task that may require the final curation/proofreading step to obtain accurate results. In that sense, the use of CartoCell crucially reduces the proofreading time from 3-5 days to just 12 ± 6 minutes per cyst (**Figure S3** and **Table S4**, **STAR Methods**).

Accounting with such a large number of samples is the key to quantifying the cell morphology patterns in these epithelial structures. We uncover two different “cell morphology” patterns within the cysts (**Figure 2**). First, in the case of the "cell height", “cell basal surface”, “cell volume” and “cell surface area” features, the cells present a clear increase in the values from top to lateral bottom. Then, cells with lower values also appear in the bottom-center of the cyst (**Figure 2D, 2E** and **Figure S5**). A different pattern can be easily distinguished when comparing the polar histograms of those characteristics, with that of “cell apical area” (comparison between **Figure 2E** and **Figure 2G**). In this second case, cells with a larger apical area are distributed at the lateral top and lateral bottom of the cysts. Importantly, we also found features that do not show any spatial pattern on the cyst (**Figure S5**), suggesting that different geometric features are independent of others. Our approach also reveals that the complexity of the cyst can reach even the level of the packing and connectivity of the cells. Here we show the example of the scutoids. Although the 3D maps of the presence of scutoids do not reveal any clear pattern (**Figure 2K**), the power of the high-content approach reveals a clear accumulation of scutoids on the bottom side of the cysts when compared with non-scutoidal cells (**Figure 2M**). Furthermore, our findings demonstrate that changes in the microenvironment, such as hypoxia, have the potential to alter cell morphology patterns (**Figure 3C, 3D**, **Figure S6** and **Table S6**).

At the tissue level, we show that cysts can adopt a variable range of shapes beyond being symmetric spheres ^56,74^ (**Figure 2A, 2B**, **Figure S2** and **Table S1**). This finding reveals a degree of complexity of the MDCK cysts that allows us to study in detail the interplay between the shape of the whole structure and the individual cell morphology. Indeed, we find that in 10-day cysts (when the size of the cyst and the number of cells increase) it is possible to find a correlation between some cellular geometric patterns and the shape of the cysts (**Figure 2H-J**, **Figure S6** and **Table S5**). Our Single-cell Cartography methodology demonstrates that the first hints of asymmetry can emerge even in tissues where there is no cell differentiation. Essentially, our method sheds light on a very basic degree of variation at both cell and global level, that was not deeply described before in cyst cultures. Taken together, our results reinforce the usefulness of this simple system to study 3D morphogenesis and help to answer complex questions such as “How do cells with different characteristics self-organize themselves in a tridimensional epithelial tissue?” or “Are cells with different shapes physiologically equivalent?”. In addition, the extracted morphological and connectivity cell information could be used to feed biophysical models and force inference analysis^10,12,75^ providing valuable knowledge of cell mechanics.

Our high-content analysis presents several advantages related to the versatility and efficiency of the method. By working with low-resolution images, it saves time during image acquisition by allowing the capture of several samples in parallel. Moreover, it accelerates image processing (segmentation and feature extraction), while also minimizing the storage space needed, and reducing photobleaching effects on the samples opening the possibility of applying CartoCell to *in vivo* imaged epithelia to study tissue development and its dynamical events. In fact, our pipeline could be combined with cell lineage analysis methods that use nuclei segmentation and tracking to study how different cell lineages remodel their geometric and connectivity features during tissue development ^34,35^. For other epithelial systems or other cell membrane markers, the strategy that we present here can be adapted. In this case, to automatically segment a large number of low-resolution samples, the users should obtain a new dataset of images and follow our pipeline (Phase 1 to Phase 5, **Figure 1**). Following that premise, we challenged our method by acquiring a large dataset of mouse embryoids and applying the whole CartoCell segmentation pipeline on it (**Figure 3F**). Remarkably, although the embryoids presented very heterogeneous morphologies, we achieved segmentation metrics comparable to those observed in the MDCK cysts (**Table S3**). Finally, CartoCell is also able to obtain accurate results in epithelial organs as the *Drosophila* egg chamber, even when the cell membranes were labeled with a sole membrane marker (**Figure 3G** and **Table S3**). The adaptability and generalizability of our approach could offer potential solutions to overcome several challenges in the field of organoids or tissue engineering ^76,77^.

CartoCell and the Single-cell Cartography methodology can unveil hidden patterns in a simple and visual manner, which is pivotal to improve the study of the self-organization of complex epithelial tissues where cells are in close contact with each other ^57,58,75,78,79^. In a biomedical context, the possibility of analyzing a very large number of samples is ideal for testing the reproducibility of epithelial organoids cultures and performing detailed comparisons between physiological and pathological conditions. Furthermore, high-throughput drug testing on animal or human epithelial organoids could take advantage of our approach to automatically analyze the effect of every drug against a target disease at the cellular level ^56,80,81^.

## LIMITATIONS OF STUDY

While the strategy presented in this work shows promise for being adapted to various epithelial systems or alternative cell membrane markers, its applicability to untested epithelial models cannot be assured. The lack of public datasets has restricted the scope of the testing to our own epithelial models. Additionally, certain factors could limit the application of this approach. The feature values related to cell connectivity could not be very feasible without careful manual proofreading to ensure accuracy. Furthermore, the effectiveness of the method may be reduced when dealing with intricate epithelial tissues featuring multiple layers or cells with complex shapes, such as pseudostratified epithelia. Despite these limitations, the strategy’s versatility remains evident, and its potential contribution to the analysis of epithelial tissues is noteworthy.

## ACKNOWLEDGMENTS

This work is funded by the Spanish Ministry of Science and Innovation Ministry of Science (PID2019-103900GB-I00) and Programa Operativo FEDER Andalucía 2014-2020 (US-1380953) to L.M.E.. L.M.E. and J.A.A.-S. work has been funded by the Junta de Andalucía (Consejería de economía, conocimiento, empresas y Universidad) grant PY18-631 co-funded by FEDER funds. A.T. has been funded by a “Contrato predoctoral PIF” from Universidad de Sevilla. C.G-V has been funded by a “Contrato predoctoral para la formación de doctores” BES-2017-082306. G.B. was supported by a Comunidad de Madrid contract (CAM), and a FPI grant from MINECO (BES-2022-077789). F.M.-B. was supported by MICINN (PID2020-120367GB-I00), and Fundación Ramón Areces (CIVP18A3904). P.G-G has been funded by Margarita Salas Fellowship - NextGenerationEU. C.H.F.-E has been funded by María Zambrano Fellowship - NextGenerationEU. I.A-C. would like to acknowledge that his work has been partially supported by the University of the Basque Country UPV/EHU grant GIU19/027, and by the Ministerio de Ciencia, Innovación y Universidades, AEI, under grant PID2021-126701OB-I00. L.M.E. also wants to thank PIE-202120E047-Conexiones-Life network for networking and input.

## AUTHOR CONTRIBUTIONS

J.A.A.-S.R. and D.F.-B. carried out all the computational experiments. D.F.-B. and I.A.-C. implemented the 3D ResU-Net network architecture. C.G.-V., L.M.-C., C.H.F.-E., and G.B. conducted the biological experiments. V.A., M.P.G. and F.M.-B. supervised cell culture experiments. C.G.-V., L.M.-C., C.H.F.-E, J.A.A.-S.R. and P.G.-G. segmented and curated the MDCK cysts, mouse embryoids and *Drosophila* egg chambers confocal images. L.M.E., I.A.-C., P.G.-G., J.A.A.-S.R., C.G.-V. and D.F.-B. designed the figures. L.M.E., I.A.-C. and P.G.-G. thought up the study, supervised the experiments and wrote the paper with input from all authors. All authors participated in the interpretation of the results, discussions, and the development of the project.

## DECLARATION OF INTERESTS

The authors declare no competing interests.

## STAR METHODS

### RESOURCES AVAILABILITY

#### Lead contact

Further information and requests for resources and reagents should be directed to and will be fulfilled by the lead contact, Luis M. Escudero.

#### Materials availability

No new materials were generated in this study.

#### Data and code availability

All data used in our analysis has been deposited at Mendeley Data and are publicly available as of the date of publication. DOIs are listed in the key resources table. All original code used in our analysis has been deposited at Mendeley Data and is publicly available as of the date of publication. DOIs are listed in the key resources table. Any additional information required to reanalyze the data reported in this paper is available from the lead contact upon request.

### EXPERIMENTAL MODEL AND STUDY PARTICIPANT DETAILS

#### MDCK cyst cell culture

Type II MDCK (Madin-Darby canine kidney) cells were maintained in minimum essential medium (MEM) containing GlutaMAX (Gibco) and supplemented with 10% fetal bovine serum (FBS), 100 U/ml penicillin and 100 µg/ml streptomycin, in a 5% CO_2_ humidified incubator at 37°C. For cyst formation, MDCK cells (2500 cells/well) were suspended in a complete medium containing 2% Matrigel (Corning, Life Sciences). Cell suspension was plated in a 4-well culture slide (Corning, Life Science) on a thin layer coating of 100% Matrigel. The plates were kept at 37°C in a humidified atmosphere of 5% CO_2_ for 4, 7 or 10 days and the medium was changed every 2 days.

The cysts under hypoxia conditions ([O_2_] = 1%) were maintained in incubators that allowed a precise and stable control of temperature, as well as O_2_ and CO_2_ concentrations. The exposure times to hypoxia were 4, 7, or 10 days, and the medium was changed every 2 days.

#### *Drosophila* egg chambers

For *Drosophila* ovary *in vivo* model, we used Resille-GFP (II) transgenic flies expressing a membrane marker tagged with the green fluorescence protein (GFP) ubiquitously to visualize the follicle cells of the egg chambers that compose the ovaries ^82^. *Drosophila* egg chambers progress through 14 morphologically distinct stages of development. For *in vivo* acquisition, 1-2-day-old females were kept in a new food vial with yeast for optimal ovary development for 48 hours. The ovaries were dissected in M3 insect culture medium (S83981L, Sigma) supplemented with 10% FBS (10270106; Gibco) and 0,20 mg/ml insulin (I550050G; Sigma). Ovarioles were isolated by removing the muscle sheath covering the ovary to avoid contraction movements during image acquisition. The ovarioles from different ovaries were mounted in a glass-based dish (Thermo Fisher Scientific), covered with supplemented medium to prevent the evaporation during *in vivo* acquisition. We obtained and processed confocal stacks from stage 2 to stage 7 egg chambers (20 egg chambers analyzed, **Table S7**). It should be noted that we distinguished between the early stages 2-3 (with an average cell count of 107.6 ± 31.5), middle stages 4-5 (315.5 ± 103.6 cells/egg chamber) and late stages 6-7 (853.8 ± 126.2 cells/egg chamber) (**Table S7**).

#### Mouse embryoids

##### mES cell derivation

For WT mouse embryonic stem (mES) cell derivation, eight-cell stage mouse embryos were recovered from the oviducts of pregnant females and cultured in KSOM (MR-020P-5F, Millipore) containing the inhibitors 2i/LIF to preserve naive pluripotency: 1μM MEK inhibitor PD0325901 (72182, STEMCELL Technologies), 3μM GSK3 inhibitor CHIR99021 (72052, STEMCELL Technologies) and 10 ngml−1 LIF (Qk019, Qkine). After 24h, the medium was changed to N2B27 medium with 2i/LIF for 48h. N2B27 medium was comprised of a 1:1 mix of DMEM F12 (21331-020, Thermo Fisher Scientific) and neurobasal A (10888-022, Thermo Fisher Scientific) supplemented with 1% v/v B27 (10889-038, Thermo Fisher Scientific), 0.5% v/v N2 (homemade), 100μM β-mercaptoethanol (31350-010, Thermo Fisher Scientific), penicillin–streptomycin (15140122, Thermo Fisher Scientific) and GlutaMAX (35050061, Thermo Fisher Scientific). Hatched blastocysts were next plated on mitomycin C-treated MEFs in Fc medium containing DMEM (41966, Thermo Fisher Scientific), 15% FBS (Stem Cell Institute), penicillin–streptomycin (15140122, Thermo Fisher Scientific), GlutaMAX (35050061, Thermo Fisher Scientific), MEM non-essential amino acids (11140035, Thermo Fisher Scientific), sodium pyruvate (11360070, Thermo Fisher Scientific) and 100μM β-mercaptoethanol (31350-010, Thermo Fisher Scientific. 2i/LIF was also added to Fc medium. Two days later, blastocysts outgrowths were trypsinized and plated to obtain mES cell colonies.

##### mES cell culture

mES cells were routinely cultured in gelatin-coated plates in Fc medium supplemented with 2i/LIF at 37°C, 5% CO_2_, 21% O_2_. Cells were routinely tested for mycoplasma contamination. For embryoid formation, mES cells (20,000 cells/well) were suspended in a complete medium containing 5% Matrigel (Corning, Life Sciences). Cell suspension was plated in an 8-well culture slide (Corning, Life Science) on a thin layer coating of 100% Matrigel. The plates were kept at 37°C in a humidified atmosphere of 5% CO_2_ for 72h and the medium was changed every 2 days.

#### Immunostaining and confocal imaging

Cysts grown on 4-well chamber slides (Corning, Life Science) were fixed with 4% paraformaldehyde in PBS and permeabilized with 0.5% TritonX-100 in Dubelcco’s Phosphate Buffered Saline (DPBS, Sigma-Aldrich) for 15 min at room temperature (RT). After blocking with a solution of 0.02% Saponin (Sigma) and 3% BSA (Applichem) in DPBS for 2h at RT, cysts were incubated overnight at 4°C with anti-β-catenin antibody (1:1000 in DPBS-0.02% Saponin-3% BSA; rabbit, Sigma-Aldrich). The following day, the cysts were washed with DPBS-0.02% Saponin-3% BSA solution (3x, 5 min each) and incubated for 90 min at RT in this solution plus anti-rabbit conjugated to Alexa Fluor 488 (1:800, Thermo Fisher Scientific) and phalloidin-Alexa-Fluor 647 (0.08 µM, Thermo Fisher Scientific). After washing with DPBS, plastic chambers were removed from microscope slides and coverslips were mounted onto slides using Fluoromount-G mounting solution (SouthernBiotech).

Cysts were imaged using a Nikon Eclipse Ti-E laser scanning confocal microscope. High-resolution Z-stack confocal images were captured using a dry x40 objective (0.95 NA) and variable zoom (3.5-5.5), with a step size of 0.5 µm per slice and a scan speed of 0.25 ms, from top to bottom. Then, they were exported as nd2 files with an XY resolution ranging between 0.06-0.08 µm per pixel and an image size of 1024 × 1024 pixels. Low resolution images were captured (from top to bottom) using x20 oil objective (0.75 NA), step size of 0.7 µm per slice, scan speed of 0.5 ms and exported as nd2 files with an XY resolution of 0.62 µm per pixel and an image size of 1024 × 1024 pixels.

*Drosophila* egg chambers, images were acquired using a Leica Stellaris 8 FALCON Confocal microscope at 25°C, with x25 water objective (0.95 NA). The whole ovarioles were captured with a resolution of (0.61 µm per pixel in XY and 1.02 µm between Z slices, using the optimal z-step size. Laser compensation was applied to maintain similar levels of GFP intensity along the z-axis.

Embryoids grown on 8-well chamber slides (Corning, Life Science) were fixed with 4% paraformaldehyde in PBS and permeabilized with PBS + 0.2% Triton Tx-100 + 0.2% SDS for 10 min at RT. After blocking with a solution of PBS + 3% BSA for 2h at RT, cysts were incubated overnight at 4°C with anti-β-catenin antibody (1:1000 in 3% BSA; rabbit, Santacruz). The following day, the cysts were washed three times with PBS and incubated for 120 min at RT in PBS + 3% BSA with Alexa-Fluor 488 anti-Rabbit (1:1000, Thermo Fisher Scientific), Alexa Fluor 555 Phalloidin (1:1000, Thermo Fisher Scientific) and DAPI (1:2000, Sigma-Aldrich). Cysts were imaged using a Nikon Eclipse Ti-E laser scanning confocal microscope. Low resolution images were captured (from top to bottom) using x20 oil objective (0.75 NA), step size of 0.7 µm per slice, scan speed of 0.5 ms and exported as nd2 files with an XY resolution of 0.62 µm per pixel and an image size of 1024 × 1024 pixels.

### METHOD DETAILS

#### Custom DNN architecture: 3D ResU-Net

Building upon the state of the art, we have designed 3D ResU-Net, a stable 3D residual U-Net ^63^ to segment epithelial cysts at the cell level. The architecture is presented in **Figure S1**. More specifically, 3D ResU-Net is formed by full pre-activation residual blocks (two 3 × 3 convolutional layers with a shortcut as shown in **Figure S1**), with 52 filters in the first level and adding 16 more at each level, dropout of 0.1 at each block. Down-sampling (max-pooling) operators are performed only in 2D, since the input volumes are anisotropic. The total number of trainable parameters is 1.3M.

The network received raw cyst images as input and outputs three different channels: i) *cell masks*, with the probability that a voxel belongs to an individual cell, ii) *contour*, containing the probabilities of cell outlines, and iii) *cell region*, representing the foreground probability of the complete cyst.

#### Network optimization

To find the best solutions with our custom 3D ResU-Net, we made an exhaustive search of hyperparameters (**Table S2**) and training configurations, exploring different loss functions, optimizers, learning rates, batch sizes, and data augmentation techniques. In particular, we minimized the binary cross-entropy (BCE) loss using the Adam optimizer, with a learning rate of 0.0001, a batch size value of 2 and using a patch size of 80×80×80 voxels. We used a Tesla P40 GPU card to train the network until convergence, i.e., for 1300 epochs with a patience established at 50 epochs monitoring the validation loss and picking up the model that performs best in the validation set (2 samples of “*training high-resolution dataset”* were used for model M1 and model M2 validation). Moreover, we applied on-the-fly data augmentation with random rotations, vertical, horizontal and Z-axis flips and brightness distortions.

#### Image preprocessing

Before accessing the network, all raw images were preprocessed for contrast homogenization using Fiji ^62^ macros. In this preprocessing, a contrast adjustment was performed using the ’enhance contrast’ function with 0.3% of saturated pixels. Additionally, an 8-bit transformation is applied to them.

In the case of high-resolution images, a down-sampling was applied using Fiji macros, transforming the variable images resolutions (0.06-0.08 µm per pixel in XY and 0.5 µm between Z slices) to have the pixel size of the low-resolution images (0.62 µm per pixel in XY and 0.7 µm between Z slices). This was done by calculating a correction factor that multiplies the size of the original image:

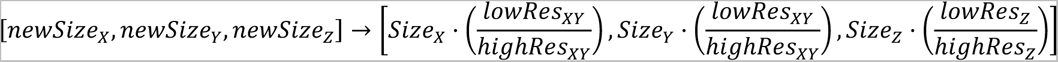

Both preprocessing Fiji macros are available at the public repository of the laboratory (see **Data and code availability** section). Note that for label down-sampling the resize command “Size” must have the interpolation method set as “None” and the option of “average when down-sampling” disabled. For raw images, however, the interpolation method should be set as “Bilinear” and the option of “average when down-sampling” ticked. In the case of the low-resolution images, before applying the aforementioned automatic contrast enhancement process, a manual preprocessing was performed in which each cyst was cropped. This procedure was carried out with Fiji by drawing the region of interest (ROI) using the "rectangle" tool and cropping using the "Crop" or "Duplicate" command.

#### Inference and postprocessing

Each patch of 80×80×80 voxels was processed by the network to reconstruct the output to the original cyst image size applying a padding of 16x16x16. In this way, we avoided border effects on every patch, as reported in ^63^. The DNN produced three different outputs representing the probabilities of the individual cell masks, contours and the whole cell region. We binarized the first two outputs based on a fixed set of threshold values (0.2 was experimentally found to work best) and created instance seeds to be fed to a marker-controlled watershed algorithm.

As a final post-processing step, we ran the Voronoi algorithm along the three dimensions, using the resulting instances of the watershed algorithm as Voronoi seeds, and the cell region output (binarized using Otsu thresholding ^83^, as the bounded region to be occupied by Voronoi cells (**Figure S3**). Thus, the unoccupied intercellular space of the cell region was filled by the nearest individual cells (Voronoi seeds).

#### Training and test datasets acquisition

*“High-resolution label images”* were obtained after segmentation of the “*high-resolution raw images”* (21 cysts) using LimeSeg ^61^, a plugin of Fiji ^62^ for 3D segmentation, based on surface elements (“Surfels”). This software was sourced from a set of seeds, manually placed over the volumetric image to localize every single cell. These seeds grow until the cell outlines are identified by detection of intensity gradient changes. The output of LimeSeg was processed using an in-house Matlab program (2021a MathWorks) to detect and curate imperfections during cysts segmentation (see **Proofreading of segmented cysts** section). A down-sampled version of these 21 segmented cysts (see **Images preprocessing** section) was used for training the model M1 (Phase 2, **Figure 1**): 19 cysts composing the training dataset, and the remaining 2, making up the validation dataset.

After running by default our high-content pipeline, we used the trained model M2 (Phase 5, **Figure 1**) to infer and subsequently segment the *“test raw images”* (60 cysts acquired at low resolution). These segmented images were manually curated using our in-house Matlab proofreading program, obtaining the *“test label images”*.

#### Proofreading of segmented cysts

A custom program developed in Matlab and available in the public repository of the laboratory (see **Data and code availability** section) was designed for the proofreading of segmented cysts. The software includes a user-friendly graphical user interface (GUI) that allows to remove, modify, merge or create labels by drawing on the two-dimensional slices of the image stack, also allowing the interpolation between labels on different slices for faster curations. Both cell and lumen labels can be modified using the GUI, which also has specific tools for each of them to ensure a proper visualization. In view that our biological study was developed on single-layer and single-lumen cysts, the proofreading software relies on a segmentation error detection tool specific to our purpose. The GUI displays the cell IDs of cells that do not contact the apical and/or basal surface of the cyst.

The software was designed to work quickly on batches of cysts. Once the cyst stops displaying errors in the GUI and is marked by the user as fixed, the next cyst will be displayed in the GUI to be corrected.

The procedure we carried out for the cyst curation started with the creation of 3 folders: One of them containing the batch of labels predicted by our pipeline, another one, the batch of raw images and a third one reserved for the curated labels. The software merges the raw images and the labels, and displays the result in a GUI along with information on possible segmentation errors. An expert reviewed the displayed image by carefully comparing each label with the staining of the cell membrane of the raw image, and adjusting the labels until a perfect segmentation was achieved.

## QUANTIFICATION AND STATISTICAL ANALYSIS

### Comparison with the state of the art

Three state-of-the-art methods were tested against our protocol, being these methods PlantSeg ^23^, Cellpose (Stringer et al., 2021) and StarDist 3D ^25^. For a more robust comparison, each one of the methods were trained 10 times using default configuration values, and the 21 low-resolution cysts used to train our model M1 (Phase 2, **Figure 1**) were inputted as training dataset. Each of the trained models was evaluated on the same test set, composed by 60 perfectly annotated cysts, to obtain measures of the error in the results yielding the table (**Table S3**).

The training of Cellpose was conducted locally by using a GPU (Graphics Processing Unit) and not using a pretrained model, as per the instructions provided in the training documentation (https://cellpose.readthedocs.io/en/latest/train.html). Inference was performed by following the instructions given in the command line documentation (https://cellpose.readthedocs.io/en/latest/command.html) and using the diameter suggested by the Cellpose GUI.

StarDist 3D training was performed in Google Colab using the official ZeroCostDL4Mic ^72^ implementation (https://github.com/HenriquesLab/ZeroCostDL4Mic/wiki) using default values except for the following parameters: patch size, which was changed to 48 and patch height, which was changed to 32 for convenience given the size of the images to be used.

PlantSeg training was performed locally following the training documentation instructions (https://github.com/hci-unihd/plant-seg) using as default configuration the 3D U-Net example for confocal imaging (https://github.com/wolny/pytorch-3dunet/blob/master/resources/3DUnet_confocal_boundary/train_config.yml) replacing patch size to [32, 64, 64] as [z, x, y] for convenience given the size of the images used and the minimum values allowed by PlantSeg. Further to the training of the network, the PlantSeg GUI has modifiable parameters for the postprocessing. This part was performed in a custom way trying to optimize the watershed (done with Simple ITK) output obtained from the probability maps predicted by the network. The parameters used were: Under-/Over-Segmentation Factor=0.75, Run Watershed 2D=False, CNN Prediction Threshold=0.113, Watershed Seeds Sigma=1.0, Watershed Boundary Sigma=0.4, Superpixels Minimum Size=1, Cell Minimum Size=5.

These three methods with identical training configurations as previously described, have also been evaluated after training them with the same 314 cysts used for training M2 model (Phase 4, **Figure 1**). To ensure a robust comparison, ten models were trained for each method.

### Cyst features and shape classification

Using an in-house Matlab code, we quantified a set of geometrical and topological parameters of the segmented epithelial cysts (**Table S1**) as is graphically described in **Figure S4**. We carried out a classification of cysts depending on the morphology and differences between axes lengths (**Figure 2A**). We considered that two axes lengths were different if they differed more than 10%. We classified all cysts into 5 groups: 1. Sphere, when the lengths of the three axes of symmetry were similar. 2. Oblate, when two axes lengths were similar and the different one was the shortest axis length. 3. Prolate, when two axes lengths were similar and the different one was the longest axis length. 4. Ellipsoid, when the three axes lengths were different. 5. Negative curvature, when the solidity (volume / convex volume) of the cyst was inferior to 0.9.

### Error evaluation of biological features

We extracted the features of both manually curated cysts (ground-truth) and the output of our high-content segmentation pipeline (without proofreading). Some of the segmented cysts without curation presented under-segmentation that promoted gaps in the segmented tissue. This defective segmentation was called "cyst opening". These gaps prevented the identification of the lumen of the “open cysts” automatically, and thus some biological features could not be extracted. The 13% (46 cysts) of the automatically segmented cysts presented this defect, and they were not used in the comparison of the biological features values (**Figure S3**). For the remaining 87% of cysts (307 cysts), features were automatically extracted and compared with the features extracted from manually curated cysts.

For each cyst, measurements of every feature were compared by computing the relative error calculated as 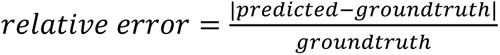. case of percentage of scutoids, we could not calculate the relative error because in some cases this feature represented a 0%, resulting in indetermination. Therefore, for the calculation of its relative error, we defined the complementary of this feature (100% - percentage of scutoids) such that we did not find any cyst with the 100% of cells being scutoids. Finally, we calculated the mean and standard deviation of the errors for every feature (**Table S4**).

### Single-cell Cartography representation

We performed an analysis of the spatial distribution of features from more than 20,000 cells from 353 segmented cysts using our Single-cell Cartography tools available in the public repository of the laboratory (see **Data and code availability** section). Different types of representations arose from the use of these tools:

*Computer rendering of 3D cysts* (**Figure 2D, 2F, 2K**, **Figure 3C** and **Figure S6**). We displayed a 3D visualization of the segmented cysts and, using a gradient of color over the cell surfaces. We can plot the normalized value of individual cell features, using our custom Matlab function Paint3D. A batch processing of cysts allowed the creation of large sheets with the previously described three-dimensional representations of cysts on which to perform a visual pattern analysis.

*Polar plots* (**Figure 2E, 2G**, **Figure 3D**, **Figure S5**, and **Figure S6**). We used two types of two-dimensional polar plots. Polar scatter plots and polar histograms were used to represent the relative spatial position (**Figure 2C**) and frequencies of a normalized cell feature for all cells of all cysts simultaneously (**Figure 2E, 2G**, **Figure 3D**, **Figure S5** and **Figure S6**). For the creation of these two-dimensional polar plots (both polar scatter plot and polar histograms) we proceeded as follows: The polar coordinate center for each cyst was set at the centroid of the cyst. The radius was normalized from 0 to 1, being 1 the distance to the farthest cell centroid from the centroid of the cyst. The colatitude angle (representing the height on the vertical axis) was calculated with respect to the horizontal plane passing through the cyst at the centroid, thus having positive angles for cells above the cyst centroid and negative angles for cells below the cyst (**Figure 2C**). The azimuthal angle (which rotates around the vertical axis) was ignored since the scope of the study was to search for patterns along the vertical axis. Disregarding this angle led to a two-dimensional representation. This approach consisted of 5 polar scatter plots and 4 polar histogram plots. First, a general polar scatter plot was shown in which all cells were represented (**Figure 2E, 2G**, **Figure 3D**, **Figure S5** and **Figure S6**). The value of the features was represented by a color gradient as in the previous case. The rest of the plots were dedicated to different ranges of the normalized feature to be studied: 0-0.25, 0.25-0.50, 0.50-0.75, 0.75-1. Each of the ranges was analyzed with a polar scatter plot and a polar histogram plot showing, normalized, the distribution of cells along the colatitude. In this way, we were able to visualize all the values of a particular cell feature distributed along the cysts vertical axis.

*Normalized cell spatial data* (**Figure 2H, 2I, 2L, 2M** and **Figure S6**). For each cell, the following data were represented: the Z-position of the cell centroid with respect to the centroid of the lowest cell in the cyst (**Figure 2H, 2I, 2M** and **Figure S6**); the distance from the cell centroid to the vertical (Z) axis passing through the centroid of the cyst (**Figure 2L** and **Figure S6**) and the value of the cellular characteristic to be studied regarding its spatial position (**Figure 2H, 2I** and **Figure S6**). The cell features were normalized regarding the maximum and minimum value of the feature in the whole cyst.

### Evaluation metrics

To evaluate our results, we used common metrics to measure instance segmentation performance in 2D and 3D images, which are calculated by matching the ground-truth and prediction segmentation masks with an Intersection Over Union (IoU) value over a certain threshold. In particular, we show values that require at least 30%, 50% and 75% IoU with the ground-truth for a detection to be a true positive (TP) (**Table S3**). More specifically, we used the following metrics:

*Precision*, defined as

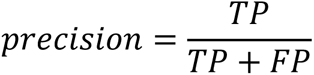

where TP and FP are the number of true and false positives, respectively.

*Recall*, defined as

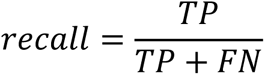

where FN is the number of false negatives.

*Accuracy*, defined as

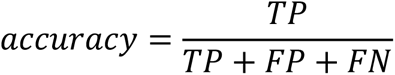

*F1 or F-score*, defined as

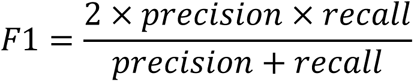

*Panoptic quality*, a unified metric to express both segmentation and recognition quality, defined as in Equation 1 of ^84^

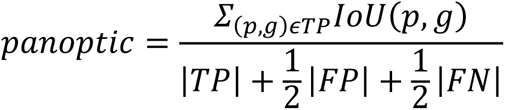

where p and g are the predicted and ground-truth segments, respectively. Therefore, 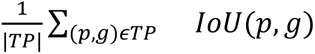 is the average of matched segments, and 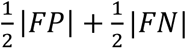 in the denominator penalizes segments without matches.

### Statistical analysis

At each time-point sampled (4 days, 7 days and 10 days cysts), at least seven independent cultures were carried out for normoxic cysts, and three independent cultures under for hypoxic cysts. For comparisons of the features values on certain cell populations between different categories of cyst shapes, as well as to compare some features between hypoxic and normoxic cysts, we used a univariate statistical protocol. First, samples were evaluated for normal distribution and similar variance by using the Shapiro-Wilk test and the two-sample F-test, respectively. If samples followed a normal distribution and similar variance, we employed the two unpaired Student’s t-test; whereas data had a normal distribution but not equal variance, we used the two-tailed Welch test. Finally, when data not adjusting to a normal distribution, we employed the non-parametric Mann-Whitney *U* test. Data were represented in a bar graph as mean ± SD (standard deviation) and p ≤ 0.05 was considered statistically significant (**Figure 2J**, **Figure 3B** and **Figure S6**). “*”, “**”, “***” and “****” indicating p ≤ 0.05, p ≤ 0.01, p ≤ 0.001and p ≤ 0.0001respectively (**Table S5** and **Table S6**). In a different statistical analysis, we tested cell spatial distribution similarity from bottom to top (in Z-axis) or from cyst centroid to outside (in XY axis) of in the proportion of scutoids and non-scutoidal cells (**Figure 2L, 2M**, **Figure S6**, **Table S5** and **Table S6**). Similarly, we also compared the similarity in the spatial distribution of cells, ranging from a colatitude angle of -90 to 90, between normoxic and hypoxic cysts regarding the proportion of cells within each range of values of the normalized feature (**Figure 2**, **Figure 3**, **Figure S5** and **Figure S6**). Following the guidelines from ^12,85^, we used the chi-square test for the trend across all samples to determine if there is a linear trend for the proportional data, considering statistically significant p ≤ 0.05 (**Table S5** and **Table S6**). “*”, “**”, “***” and “****” indicating p ≤ 0.05, p ≤ 0.01, p ≤ 0.001and p ≤ 0.0001, respectively. Statistical analyzes and graphs were performed using GraphPad Prism version 8.4.2. (GraphPad Software, La Jolla California, USA, www.graphpad.com).

## EXCEL TABLE TITLES AND LEGENDS

**Table S1. Extracted features from 353 curated cysts (104 cysts at 4 days, 103 cysts at 7 days, 116 cysts at 10 days), Related to Figure 2**. Tab1 (GlobalFeatures) shows the mean of geometric and packing features describing the whole cyst. Tab 2 (CellParameters) shows the mean and standard deviation of cellular geometric and packing features.

**Table S2. Hyperparameter search space for our proposed 3D ResU-Net, Related to Figure 1**.

**Table S3. Performance evaluation of our pipeline (CartoCell) on images of different epithelial tissues and comparison with other state-of-the-art segmentation methods, using the evaluation metrics described in (STAR Methods), Related to Figure 1**.

**Table S4. Relative error between features extracted using automatically segmented cysts and manually curated cysts (STAR Methods), Related to Figure 1**.

**Table S5. Cyst morphology and scutoids location statistics, Related to Figure 2**. Tab 1 (Cyst morphology properties) shows the statistical differences in cell proportions after comparing different cyst morphologies. Tab 2 (Stats for position of scutoids) shows the statistical comparison between the spatial position of scutoids and non scutoids cells.

**Table S6. Comparison of morphology and packing features of normoxic and hypoxic MDCK cysts, Related to Figure 2**. Tab 1 (Cyst morphology properties) shows the statistical differences in some morphology and packing features of normoxic and hypoxic cysts. Tab 2 (Stats for cell height), Tab 3 (Stats for cell apical area), Tab 4 (Stats for cell basal area), Tab 5 (Stats for cell volume), Tab 6 (Stats for cell surface area), Tab 7 (Stats for cell solidity), Tab 8 (Stats for cell aspect ratio) show the statistical comparison of the spatial position of the values of different cell features. Tab 9 (Stats for scutoids) shows the statistical comparison between the spatial position of scutoids and non scutoids cells.

**Table S7. Classification of the developmental stages of *Drosophila* egg chambers employed, Related to Figure 3**.

## Notes

https://data.mendeley.com/datasets/7gbkxgngpm

### Competing Interest Statement

The authors have declared no competing interest.

### Summary of Updates

Title updated; MS updated; Supplemental files updated;

## REFERENCES

Lecuit, T., and Lenne, P.-F. (2007). Cell surface mechanics and the control of cell shape, tissue patterns and morphogenesis. Nat Rev Mol Cell Biol 8, 633–644. 10.1038/nrm2222.

Davidson, L.A. (2012). Epithelial machines that shape the embryo. Trends Cell Biol 22, 82–87. 10.1016/j.tcb.2011.10.005.

Bryant, D.M., and Mostov, K.E. (2008). From cells to organs: building polarized tissue. Nat Rev Mol Cell Biol 9, 887–901. 10.1038/nrm2523.

Kriston-Vizi, J., and Flotow, H. (2017). Getting the whole picture: High content screening using three-dimensional cellular model systems and whole animal assays. Cytometry Part A 91, 152–159. 10.1002/cyto.a.22907.

Okuda, S., and Sato, K. (2022). Polarized interfacial tension induces collective migration of cells, as a cluster, in a 3D tissue. Biophys J 121, 1856–1867. 10.1016/J.BPJ.2022.04.018.

Okuda, S., Miura, T., Inoue, Y., Adachi, T., and Eiraku, M. (2018). Combining Turing and 3D vertex models reproduces autonomous multicellular morphogenesis with undulation, tubulation, and branching. Scientific Reports 2018 8:1 *8*, 1–15. 10.1038/s41598-018-20678-6.

Messal, H.A., Alt, S., Ferreira, R.M.M., Gribben, C., Wang, V.M.-Y., Cotoi, C.G., Salbreux, G., and Behrens, A. (2019). Tissue curvature and apicobasal mechanical tension imbalance instruct cancer morphogenesis. Nature 566, 126–130. 10.1038/s41586-019-0891-2.

Ioannou, F., Dawi, M.A., Tetley, R.J., Mao, Y., and Muñoz, J.J. (2020). Development of a New 3D Hybrid Model for Epithelia Morphogenesis. Front Bioeng Biotechnol 8. 10.3389/fbioe.2020.00405.

Gómez, H.F., Dumond, M.S., Hodel, L., Vetter, R., and Iber, D. (2021). 3D cell neighbour dynamics in growing pseudostratified epithelia. Elife 10. 10.7554/eLife.68135.

Gómez-Gálvez, P., Anbari, S., Escudero, L.M., and Buceta, J. (2021). Mechanics and self-organization in tissue development. Semin Cell Dev Biol 120, 147–159. 10.1016/j.semcdb.2021.07.003.

Gómez-Gálvez, P., Vicente-Munuera, P., Tagua, A., Forja, C., Castro, A.M., Letrán, M., Valencia-Expósito, A., Grima, C., Bermúdez-Gallardo, M., Serrano-Pérez-Higueras, Ó., et al. (2018). Scutoids are a geometrical solution to three-dimensional packing of epithelia. Nat Commun 9. 10.1038/s41467-018-05376-1.

Gómez-Gálvez, P., Vicente-Munuera, P., Anbari, S., Tagua, A., Gordillo-Vázquez, C., Andrés-San Román, J.A., Franco-Barranco, D., Palacios, A.M., Velasco, A., Capitán-Agudo, C., et al. (2022). A quantitative biophysical principle to explain the 3D cellular connectivity in curved epithelia. Cell Syst 13, 631–643.e8. 10.1016/J.CELS.2022.06.003.

Gómez-Gálvez, P., Vicente-Munuera, P., Anbari, S., Buceta, J., and Escudero, L.M. (2021). The complex three-dimensional organization of epithelial tissues. Development 148, dev195669. 10.1242/dev.195669.

Lou, Y., Rupprecht, J.-F., Hiraiwa, T., and Saunders, T.E. (2022). Curvature-induced cell rearrangements in biological tissues. bioRxiv, 2022.05.18.492428. 10.1101/2022.05.18.492428.

Prabhakara, C., Iyer, K.S., Rao, M., Saunders, T.E., and Mayor, S. (2022). Quantitative analysis of three-dimensional cell organisation and concentration profiles within curved epithelial tissues. bioRxiv, 2022.05.16.492131. 10.1101/2022.05.16.492131.

Rupprecht, J.-F., Ong, K.H., Yin, J., Huang, A., Dinh, H.-H.-Q., Singh, A.P., Zhang, S., Yu, W., and Saunders, T.E. (2017). Geometric constraints alter cell arrangements within curved epithelial tissues. Mol Biol Cell 28, 3582–3594. 10.1091/mbc.e17-01-0060.

Laine, R.F., Arganda-Carreras, I., Henriques, R., and Jacquemet, G. (2021). Avoiding a replication crisis in deep-learning-based bioimage analysis. Nat Methods 18, 1136–1144. 10.1038/s41592-021-01284-3.

Meijering, E. (2020). A bird’s-eye view of deep learning in bioimage analysis. Comput Struct Biotechnol J 18, 2312–2325. 10.1016/j.csbj.2020.08.003.

Moen, E., Bannon, D., Kudo, T., Graf, W., Covert, M., and Van Valen, D. (2019). Deep learning for cellular image analysis. Nat Methods 16, 1233–1246. 10.1038/s41592-019-0403-1.

Ciresan, D., Giusti, A., Gambardella, L., and Schmidhuber, J. (2012). Deep Neural Networks Segment Neuronal Membranes in Electron Microscopy Images. In Advances in Neural Information Processing Systems, F. Pereira, C. J. Burges, L. Bottou, and K. Q. Weinberger, eds. (Curran Associates, Inc.).

Falk, T., Mai, D., Bensch, R., Çiçek, Ö., Abdulkadir, A., Marrakchi, Y., Böhm, A., Deubner, J., Jäckel, Z., Seiwald, K., et al. (2018). U-Net: deep learning for cell counting, detection, and morphometry. Nature Methods 2018 16:1 *16*, 67–70. 10.1038/s41592-018-0261-2.

Wei, D., Lin, Z., Franco-Barranco, D., Wendt, N., Liu, X., Yin, W., Huang, X., Gupta, A., Jang, W.-D., Wang, X., et al. (2020). MitoEM Dataset: Large-Scale 3D Mitochondria Instance Segmentation from EM Images. In, pp. 66–76. 10.1007/978-3-030-59722-1_7.

Wolny, A., Cerrone, L., Vijayan, A., Tofanelli, R., Barro, A.V., Louveaux, M., Wenzl, C., Strauss, S., Wilson-Sánchez, D., Lymbouridou, R., et al. (2020). Accurate and versatile 3D segmentation of plant tissues at cellular resolution. Elife 9. 10.7554/eLife.57613.

Schmidt, U., Weigert, M., Broaddus, C., and Myers, G. (2018). Cell Detection with Star-Convex Polygons. In, pp. 265–273. 10.1007/978-3-030-00934-2_30.

Weigert, M., Schmidt, U., Haase, R., Sugawara, K., and Myers, G. (2020). Star-convex Polyhedra for 3D Object Detection and Segmentation in Microscopy. In 2020 IEEE Winter Conference on Applications of Computer Vision (WACV) (IEEE), pp. 3655–3662. 10.1109/WACV45572.2020.9093435.

Stringer, C., Wang, T., Michaelos, M., and Pachitariu, M. (2021). Cellpose: a generalist algorithm for cellular segmentation. Nat Methods 18, 100–106. 10.1038/s41592-020-01018-x.

Yan, Z., Yang, X., and Cheng, K.-T.T. (2018). A Deep Model with Shape-Preserving Loss for Gland Instance Segmentation. In, pp. 138–146. 10.1007/978-3-030-00934-2_16.

Lin, Z., Wei, D., Petkova, M.D., Wu, Y., Ahmed, Z., K, K.S., Zou, S., Wendt, N., Boulanger-Weill, J., Wang, X., et al. (2021). NucMM Dataset: 3D Neuronal Nuclei Instance Segmentation at Sub-Cubic Millimeter Scale. In, pp. 164–174. 10.1007/978-3-030-87193-2_16.

Meyer, F. (1994). Topographic distance and watershed lines. Signal Processing 38, 113–125. 10.1016/0165-1684(94)90060-4.

Cousty, J., Bertrand, G., Najman, L., and Couprie, M. (2009). Watershed Cuts: Minimum Spanning Forests and the Drop of Water Principle. IEEE Trans Pattern Anal Mach Intell 31, 1362–1374. 10.1109/TPAMI.2008.173.

Wolf, S., Pape, C., Bailoni, A., Rahaman, N., Kreshuk, A., Köthe, U., and Hamprecht, F.A. (2018). The Mutex Watershed: Efficient, Parameter-Free Image Partitioning. In, pp. 571–587. 10.1007/978-3-030-01225-0_34.

Bailoni, A., Pape, C., Hütsch, N., Wolf, S., Beier, T., Kreshuk, A., and Hamprecht, F.A. (2019). GASP, a generalized framework for agglomerative clustering of signed graphs and its application to Instance Segmentation. 10.48550/arxiv.1906.11713.

Kappes, J.H., Speth, M., Andres, B., Reinelt, G., and Schn, C. (2011). Globally Optimal Image Partitioning by Multicuts. In Lecture Notes in Computer Science (including subseries Lecture Notes in Artificial Intelligence and Lecture Notes in Bioinformatics) (Springer, Berlin, Heidelberg), pp. 31–44. 10.1007/978-3-642-23094-3_3.

He, Z., Maynard, A., Jain, A., Gerber, T., Petri, R., Lin, H.-C., Santel, M., Ly, K., Dupré, J.-S., Sidow, L., et al. (2022). Lineage recording in human cerebral organoids. Nat Methods 19, 90–99. 10.1038/s41592-021-01344-8.

de Medeiros, G., Ortiz, R., Strnad, P., Boni, A., Moos, F., Repina, N., Challet Meylan, L., Maurer, F., and Liberali, P. (2022). Multiscale light-sheet organoid imaging framework. Nat Commun 13, 4864. 10.1038/s41467-022-32465-z.

Razzak, M.I., Naz, S., and Zaib, A. (2018). Deep Learning for Medical Image Processing: Overview, Challenges and the Future. In, pp. 323–350. 10.1007/978-3-319-65981-7_12.

Perez, L., and Wang, J. (2017). The Effectiveness of Data Augmentation in Image Classification using Deep Learning. 10.48550/arxiv.1712.04621.

Shorten, C., and Khoshgoftaar, T.M. (2019). A survey on Image Data Augmentation for Deep Learning. J Big Data 6, 1–48. 10.1186/S40537-019-0197-0/FIGURES/33.

Elia, N., and Lippincott-Schwartz, J. (2009). Culturing MDCK Cells in Three Dimensions for Analyzing Intracellular Dynamics. Curr Protoc Cell Biol 43. 10.1002/0471143030.cb0422s43.

Vidal-Quadras, M., Holst, M.R., Francis, M.K., Larsson, E., Hachimi, M., Yau, W.L., Peränen, J., Martín-Belmonte, F., and Lundmark, R. (2017). Endocytic turnover of Rab8 controls cell polarization. J Cell Sci. 10.1242/jcs.195420.

Dukes, J.D., Whitley, P., and Chalmers, A.D. (2011). The MDCK variety pack: Choosing the right strain. BMC Cell Biol 12. 10.1186/1471-2121-12-43.

Martín-Belmonte, F., Yu, W., Rodríguez-Fraticelli, A.E., Ewald, A., Werb, Z., Alonso, M.A., and Mostov, K. (2008). Cell-Polarity Dynamics Controls the Mechanism of Lumen Formation in Epithelial Morphogenesis. Current Biology 18, 507–513. 10.1016/j.cub.2008.02.076.

Yonemura, S. (2014). Differential Sensitivity of Epithelial Cells to Extracellular Matrix in Polarity Establishment. PLoS One 9, e112922. 10.1371/journal.pone.0112922.

Engelberg, J.A., Datta, A., Mostov, K.E., and Hunt, C.A. (2011). MDCK Cystogenesis Driven by Cell Stabilization within Computational Analogues. PLoS Comput Biol 7, e1002030-.

Alfonso-Pérez, T., Baonza, G., Herranz, G., and Martín-Belmonte, F. (2022). Deciphering the interplay between autophagy and polarity in epithelial tubulogenesis. Semin Cell Dev Biol 131, 160–172. 10.1016/j.semcdb.2022.05.015.

Guo, Q., Xia, B., Moshiach, S., Xu, C., Jiang, Y., Chen, Y., Sun, Y., Lahti, J.M., and Zhang, X.A. (2008). The microenvironmental determinants for kidney epithelial cyst morphogenesis. Eur J Cell Biol 87, 251–266. 10.1016/j.ejcb.2007.11.004.

Herranz, G., and Martín-Belmonte, F. (2022). Cadherin-mediated adhesion takes control. EMBO J. 10.15252/embj.2022112662.

Imai, M., Furusawa, K., Mizutani, T., Kawabata, K., and Haga, H. (2015). Three-dimensional morphogenesis of MDCK cells induced by cellular contractile forces on a viscous substrate. Sci Rep 5, 14208. 10.1038/srep14208.

O’Brien, L.E., Zegers, M.M.P., and Mostov, K.E. (2002). Building epithelial architecture: Insights from three-dimensional culture models. Nat Rev Mol Cell Biol 3, 531–537. 10.1038/nrm859.

Wells, E.K., Yarborough, O., Lifton, R.P., Cantley, L.G., and Caplan, M.J. (2013). Epithelial morphogenesis of MDCK cells in three-dimensional collagen culture is modulated by interleukin-8. American Journal of Physiology-Cell Physiology 304, C966–C975. 10.1152/ajpcell.00261.2012.

Yu, W., Fang, X., Ewald, A., Wong, K., Hunt, C.A., Werb, Z., Matthay, M.A., and Mostov, K. (2007). Formation of Cysts by Alveolar Type II Cells in Three-dimensional Culture Reveals a Novel Mechanism for Epithelial Morphogenesis. Mol Biol Cell 18, 1693–1700. 10.1091/mbc.e06-11-1052.

Sakurai, A., Matsuda, M., and Kiyokawa, E. (2012). Activated Ras Protein Accelerates Cell Cycle Progression to Perturb Madin-Darby Canine Kidney Cystogenesis. Journal of Biological Chemistry 287, 31703–31711. 10.1074/jbc.M112.377804.

Schmeichel, K.L., and Bissell, M.J. (2003). Modeling tissue-specific signaling and organ function in three dimensions. J Cell Sci 116, 2377–2388. 10.1242/jcs.00503.

Yan, L., Tsujita, K., Fujita, Y., and Itoh, T. (2021). PTEN is required for the migration and invasion of Ras-transformed MDCK cells. FEBS Lett 595, 1303–1312. 10.1002/1873-3468.14053.

Fessenden, T.B., Beckham, Y., Perez-Neut, M., Ramirez-San Juan, G., Chourasia, A.H., Macleod, K.F., Oakes, P.W., and Gardel, M.L. (2018). Dia1-dependent adhesions are required by epithelial tissues to initiate invasion. Journal of Cell Biology 217, 1485–1502. 10.1083/jcb.201703145.

Beck, L.E., Lee, J., Coté, C., Dunagin, M.C., Lukonin, I., Salla, N., Chang, M.K., Hughes, A.J., Mornin, J.D., Gartner, Z.J., et al. (2022). Systematically quantifying morphological features reveals constraints on organoid phenotypes. Cell Syst 13, 547–560.e3. 10.1016/j.cels.2022.05.008.

Carleton, A.E., Duncan, M.C., and Taniguchi, K. (2022). Human epiblast lumenogenesis: From a cell aggregate to a lumenal cyst. Semin Cell Dev Biol 131, 117–123. 10.1016/j.semcdb.2022.05.009.

Metzger, J.J., Simunovic, M., and Brivanlou, A.H. (2018). Synthetic embryology: controlling geometry to model early mammalian development. Curr Opin Genet Dev 52, 86–91. 10.1016/j.gde.2018.06.006.

Voronoi, G. (1908). Nouvelles applications des paramètres continus à la théorie des formes quadratiques. Deuxième mémoire. Recherches sur les parallélloèdres primitifs. Journal für die reine und angewandte Mathematik 134, 198–287.

Honda, H. (1978). Description of cellular patterns by Dirichlet domains: The two-dimensional case. J Theor Biol 72, 523–543. 10.1016/0022-5193(78)90315-6.

Machado, S., Mercier, V., and Chiaruttini, N. (2019). LimeSeg: a coarse-grained lipid membrane simulation for 3D image segmentation. BMC Bioinformatics 20, 2. 10.1186/s12859-018-2471-0.

Schindelin, J., Arganda-Carreras, I., Frise, E., Kaynig, V., Longair, M., Pietzsch, T., Preibisch, S., Rueden, C., Saalfeld, S., Schmid, B., et al. (2012). Fiji: an open-source platform for biological-image analysis. Nat Methods *9*, 676–682. 10.1038/nmeth.2019.

Franco-Barranco, D., Muñoz-Barrutia, A., and Arganda-Carreras, I. (2022). Stable Deep Neural Network Architectures for Mitochondria Segmentation on Electron Microscopy Volumes. Neuroinformatics 20, 437–450. 10.1007/s12021-021-09556-1.

Kaliman, S., Jayachandran, C., Rehfeldt, F., and Smith, A.-S. (2016). Limits of Applicability of the Voronoi Tessellation Determined by Centers of Cell Nuclei to Epithelium Morphology. Front Physiol 7.

Raykhel, I., Moafi, F., Myllymäki, S.M., Greciano, P.G., Matlin, K.S., Moyano, J. V., Manninen, A., and Myllyharju, J. (2018). BAMBI is a novel HIF1-dependent modulator of TGFβ-mediated disruption of cell polarity in hypoxia. J Cell Sci. 10.1242/jcs.210906.

Schley, G., Scholz, H., Kraus, A., Hackenbeck, T., Klanke, B., Willam, C., Wiesener, M.S., Heinze, E., Burzlaff, N., Eckardt, K.-U., et al. (2015). Hypoxia inhibits nephrogenesis through paracrine Vegfa despite the ability to enhance tubulogenesis. Kidney Int 88, 1283– 1292. 10.1038/ki.2015.214.

Shahbazi, M.N., Scialdone, A., Skorupska, N., Weberling, A., Recher, G., Zhu, M., Jedrusik, A., Devito, L.G., Noli, L., Macaulay, I.C., et al. (2017). Pluripotent state transitions coordinate morphogenesis in mouse and human embryos. Nature 552, 239–243. 10.1038/nature24675.

Schliffka, M.F., Dumortier, J.G., Pelzer, D., Mukherjee, A., and Maître, J.-L. (2023). Inverse blebs operate as hydraulic pumps during mouse blastocyst formation. bioRxiv, 2023.05.03.539105. 10.1101/2023.05.03.539105.

Fadiga, J., and Nystul, T.G. (2019). The follicle epithelium in the Drosophila ovary is maintained by a small number of stem cells. Elife 8. 10.7554/eLife.49050.

Duhart, J.C., Parsons, T.T., and Raftery, L.A. (2017). The repertoire of epithelial morphogenesis on display: Progressive elaboration of Drosophila egg structure. Mech Dev 148, 18–39. 10.1016/j.mod.2017.04.002.

Spradling, A.C. (1993). Developmental genetics of oogenesis. The development of Drosophila melanogaster., 1–70.

von Chamier, L., Laine, R.F., Jukkala, J., Spahn, C., Krentzel, D., Nehme, E., Lerche, M., Hernández-Pérez, S., Mattila, P.K., Karinou, E., et al. (2021). Democratising deep learning for microscopy with ZeroCostDL4Mic. Nat Commun 12, 2276. 10.1038/s41467-021-22518-0.

Shrestha, P., Kuang, N., and Yu, J. (2023). Efficient end-to-end learning for cell segmentation with machine generated weak annotations. Commun Biol 6, 232. 10.1038/s42003-023-04608-5.

Cerruti, B., Puliafito, A., Shewan, A.M., Yu, W., Combes, A.N., Little, M.H., Chianale, F., Primo, L., Serini, G., Mostov, K.E., et al. (2013). Polarity, cell division, and out-of-equilibrium dynamics control the growth of epithelial structures. Journal of Cell Biology 203, 359–372. 10.1083/jcb.201305044.

Dahl-Jensen, S., and Grapin-Botton, A. (2017). The physics of organoids: a biophysical approach to understanding organogenesis. Development 144, 946–951. 10.1242/dev.143693.

Laurent, J., Blin, G., Chatelain, F., Vanneaux, V., Fuchs, A., Larghero, J., and Théry, M. (2017). Convergence of microengineering and cellular self-organization towards functional tissue manufacturing. Nat Biomed Eng 1, 939–956. 10.1038/s41551-017-0166-x.

Huch, M., Knoblich, J.A., Lutolf, M.P., and Martinez-Arias, A. (2017). The hope and the hype of organoid research. Development 144, 938–941. 10.1242/dev.150201.

Takasato, M., Er, P.X., Chiu, H.S., and Little, M.H. (2016). Generation of kidney organoids from human pluripotent stem cells. Nat Protoc 11, 1681–1692. 10.1038/nprot.2016.098.

Treacy, N.J., Clerkin, S., Davis, J.L., Kennedy, C., Miller, A.F., Saiani, A., Wychowaniec, J.K., Brougham, D.F., and Crean, J. (2023). Growth and differentiation of human induced pluripotent stem cell (hiPSC)-derived kidney organoids using fully synthetic peptide hydrogels. Bioact Mater 21, 142–156. 10.1016/j.bioactmat.2022.08.003.

Drost, J., and Clevers, H. (2018). Organoids in cancer research. Nat Rev Cancer 18, 407–418. 10.1038/s41568-018-0007-6.

Lo, Y.-H., Karlsson, K., and Kuo, C.J. (2020). Applications of organoids for cancer biology and precision medicine. Nat Cancer 1, 761–773. 10.1038/s43018-020-0102-y.

Morin, X., Daneman, R., Zavortink, M., and Chia, W. (2001). A protein trap strategy to detect GFP-tagged proteins expressed from their endogenous loci in *Drosophila*. Proceedings of the National Academy of Sciences 98, 15050–15055. 10.1073/pnas.261408198.

Otsu, N. (1979). A Threshold Selection Method from Gray-Level Histograms. IEEE Trans Syst Man Cybern 9, 62–66. 10.1109/TSMC.1979.4310076.

Kirillov, A., He, K., Girshick, R., Rother, C., and Dollar, P. (2018). Panoptic Segmentation. Proceedings of the IEEE Computer Society Conference on Computer Vision and Pattern Recognition 2019*-June*, 9396–9405. 10.48550/arxiv.1801.00868.

Sánchez-Gutiérrez, D., Tozluoglu, M., Barry, J.D., Pascual, A., Mao, Y., and Escudero, L.M. (2016). Fundamental physical cellular constraints drive self-organization of tissues. EMBO J 35, 77–88. 10.15252/embj.201592374.

