## supplemental for "CartoCell, a high-content pipeline for 3D image analysis, unveils cell morphology patterns in epithelia"

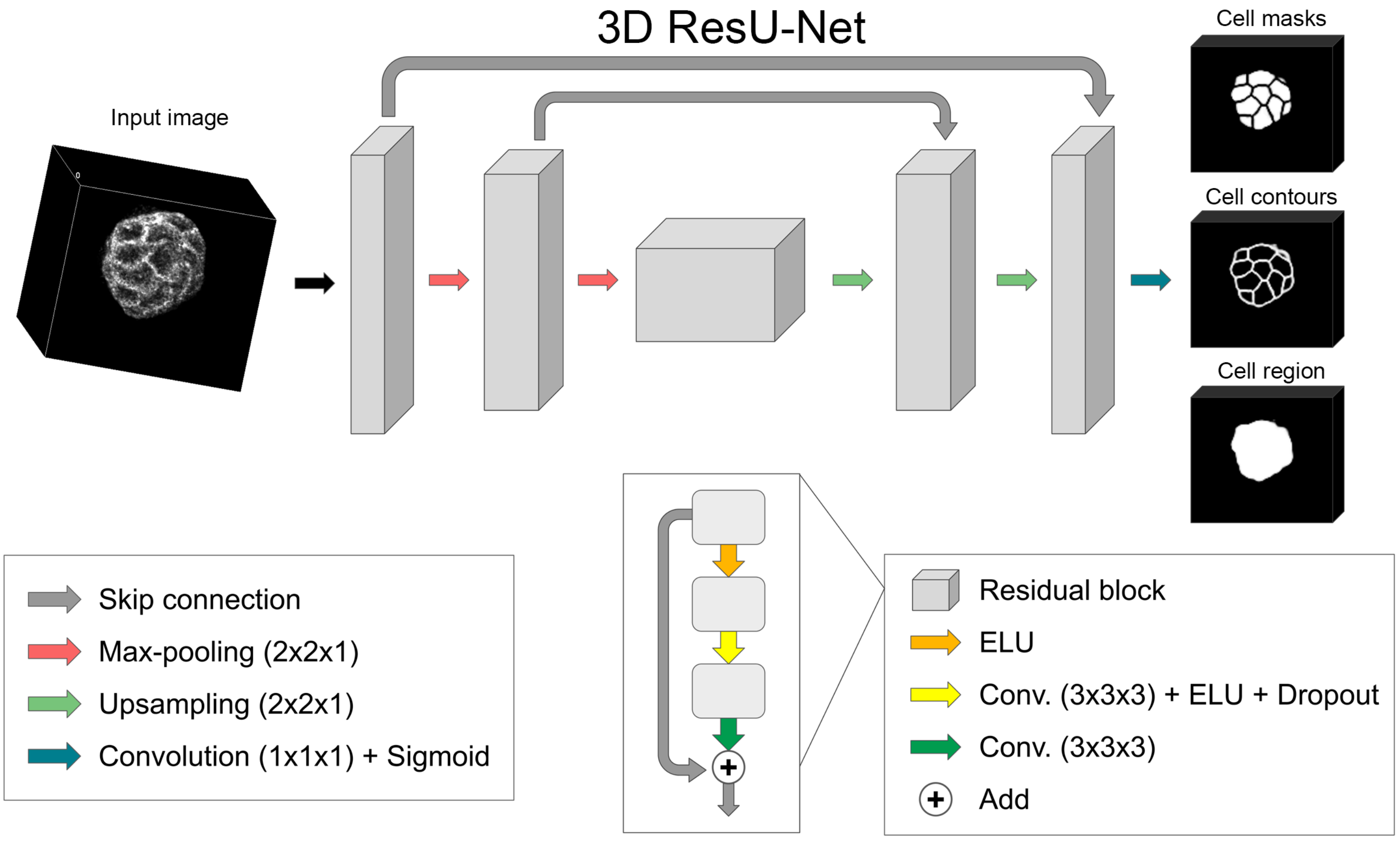
3D ResU-Net
Input image
Cell masks
Cell contours
Cell region
Skip connection
Max-pooling (2x2x1)
Upsampling (2x2x1)
Convolution (1x1x1) + Sigmoid
Residual block
ELU
Conv. (3x3x3) + ELU + Dropout
Conv. (3x3x3)
Add

**Figure S1. Architecture of the 3D ResU-Net segmentation network, Related to Figure 1.**

Our proposed 3D ResU-Net is a U-Net-like network with three depth levels and residual blocks. It receives 3D single-channel images as input and produces three different outputs representing the probabilities of the individual cell masks, contours and whole cell region. Convolutional filters are applied in 3D, while the down-sampling and up-sampling operations are only performed in 2D due to the anisotropy of image data.

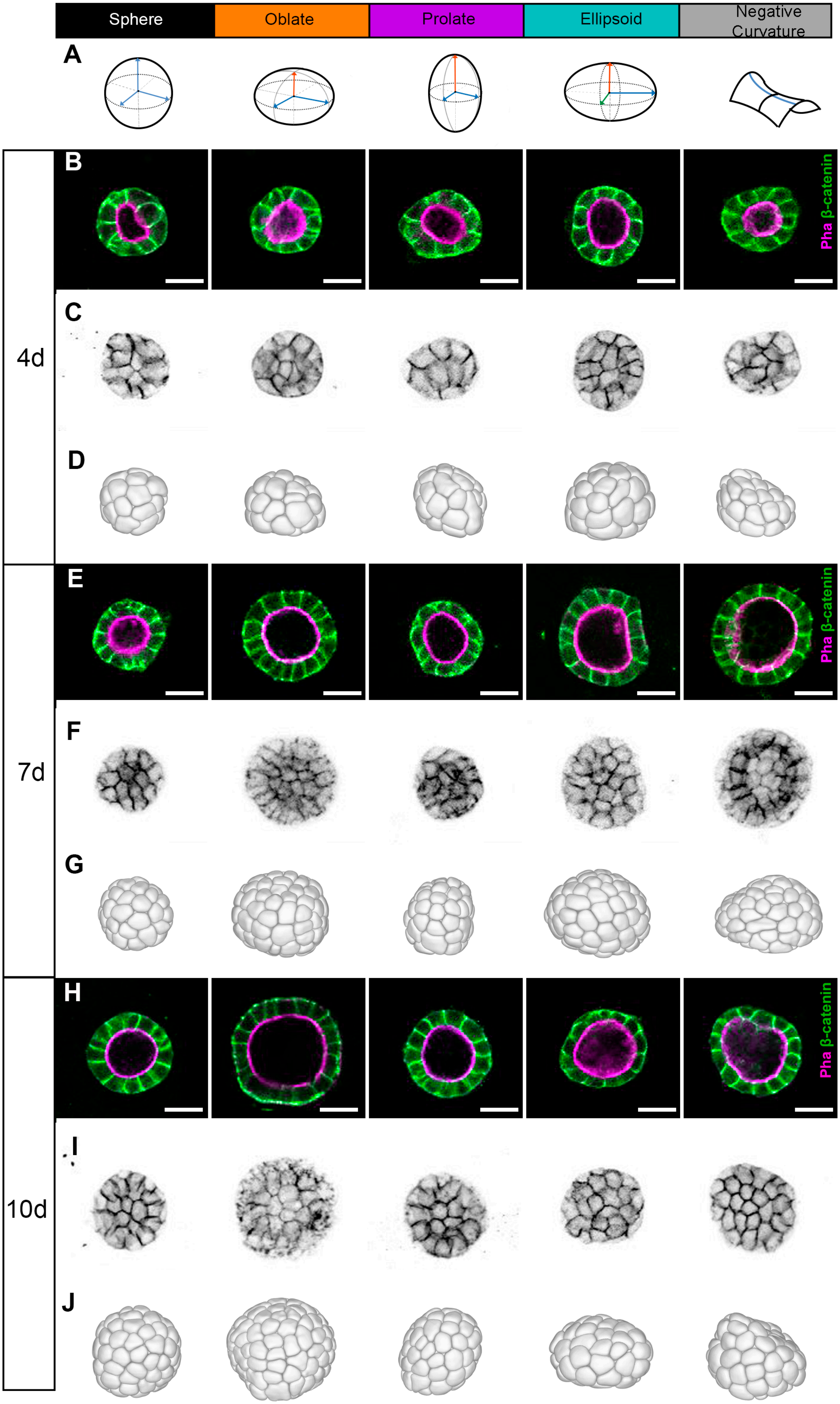
Sphere
Oblate
Prolate
Ellipsoid
Negative Curvature
A
B
Pha β-catenin
C
D
4d
E
Pha β-catenin
F
G
7d
H
Pha β-catenin
I
J
10d

**Figure S2. MDCK cysts adopt different shapes in 3D cell cultures, Related to Figure 1.**

**A**) Schematic representation of the morphological cyst classification. **B**, **E**, **H**) Middle sections of top-to-bottom confocal microscopy Z-stack images of representative cysts belonging to each morphological category at 4, 7 and 10 days. Cell contours were stained with Alexa Fluor 647 phalloidin (magenta) and anti-β-catenin antibody (green) (**STAR Methods**). Scale bars, 20 μm. **C**, **F**, **I)** Half projections of representative cysts belonging to each morphological category at 4, 7 and 10 days (cell contours stained by Alexa Fluor 647 phalloidin and anti-β-catenin antibody) (**STAR Methods**). **D**, **G**, **J**) Computer rendering (front or back view) of the segmented cysts belonging to each morphological category at 4, 7 and 10 days.

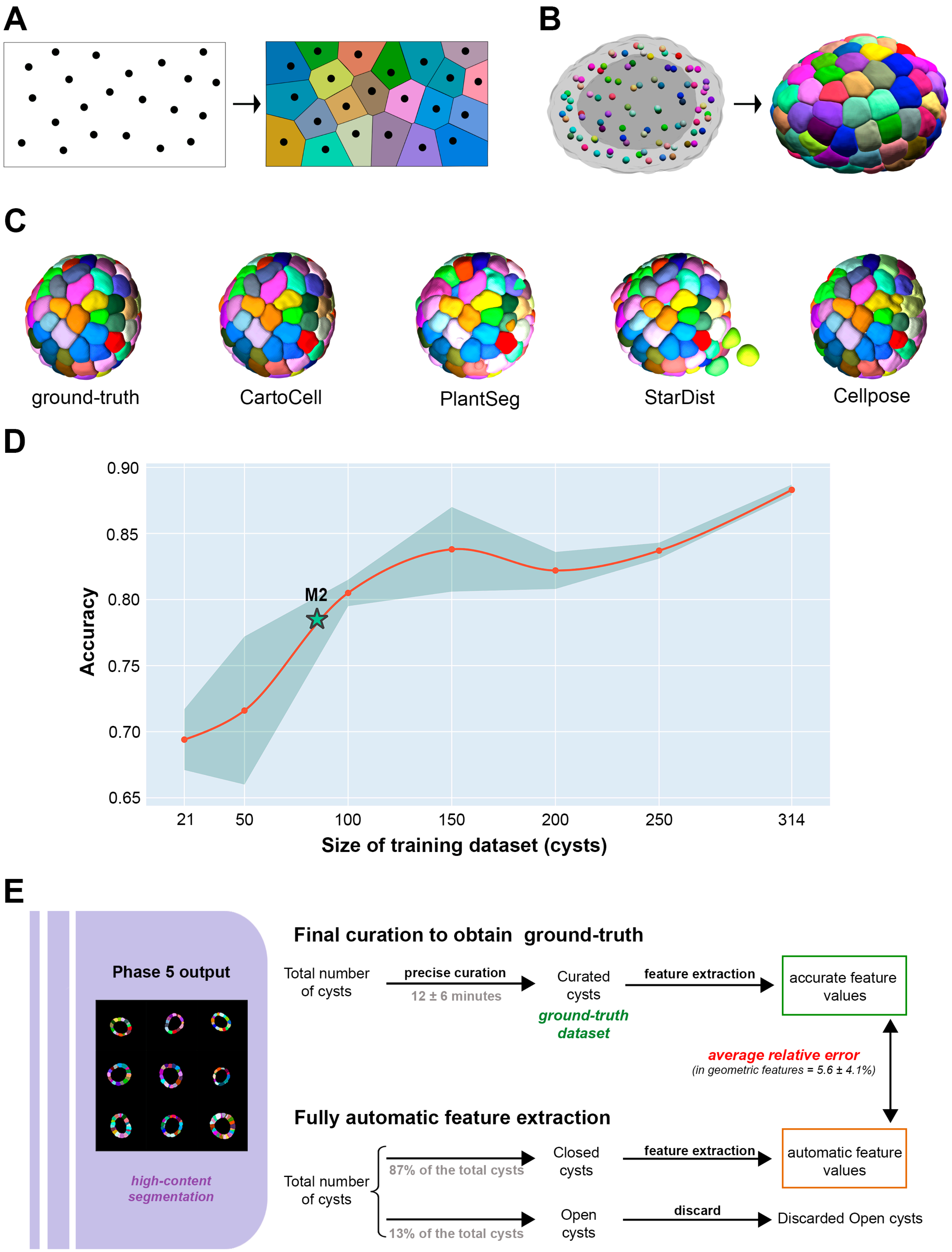
A
B
C
ground-truth
CartoCell
PlantSeg
StarDist
Cellpose
D
Accuracy
0.90
0.85
0.80
0.75
0.70
0.65
M2
21
50
100
150
200
250
314
Size of training dataset (cysts)
E
Phase 5 output
high-content segmentation
Final curation to obtain ground-truth
Total number of cysts
precise curation
12 ± 6 minutes
Curated cysts
ground-truth dataset
feature extraction
accurate feature values
average relative error
(in geometric features = 5.6 ± 4.1%)
Fully automatic feature extraction
Total number of cysts
87% of the total cysts
13% of the total cysts
Closed cysts
Open cysts
feature extraction
discard
automatic feature values
Discarded Open cysts

**Figure S3. Appendixes and details regarding Figure 1, Related to Figure 1.**

**A-C**) Voronoi tessellations and comparison of custom three-dimensional Voronoi based model against state-of-the-art methods. **A**) Voronoi tessellation in two dimensions occupying the space of a rectangle using randomly distributed seeds. **B**) Three-dimensional Voronoi tessellation of a cyst filling the space determined by the cell tissue using the centroids of each cell as seeds. **C**) Comparison of a manually labeled cyst (ground-truth) against our proposed method (CartoCell) and the three state-of-the-art methods chosen for comparison: PlantSeg, StarDist and Cellpose. **D**) Plot illustrating the relationship between the accuracy of a 3D ResU-Net model (y-axis), when trained with different number of perfectly annotated cysts (x-axis). The accuracy is measured for seven different dataset sizes: 21 (size of the dataset used to train the model M1), 50, 100, 150, 200, 250, and 314 (size of the imperfect dataset used to train the model M2). Each of these training scenarios is repeated 10 times to obtain the mean value (red dots) and standard deviation (blue band). The intersection between the accuracy trendline (red line) as the training dataset size increases, and the accuracy of the CartoCell model M2 is represented by a star symbol. This intersection indicated that around 90 fully segmented cysts are required in a single phase of training to achieve equivalent results to the whole pipeline of CartoCell with only 21 cysts as input. **E**) Appendix to Fig.1 showing two possible paths for feature extraction. After the Phase 5, a ground-truth dataset can be obtained by final manual curation of the results (12.0 ± 6 minutes per cyst) to extract the accurate feature values. On the other hand, there is an alternative option, opting for the automatic feature extraction of geometric data assuming an average relative error: Based on the data of our study, 46 of 353 cysts (13%) presented under-segmentation defects which induced open cysts and impaired the feature extraction. The extraction of the features of the remaining cyst (87%, closed cysts) provided results with an average error of 5.6 ± 4.1% in geometric characteristics compared with the accurate feature values (**Table S4** and **STAR Methods**).

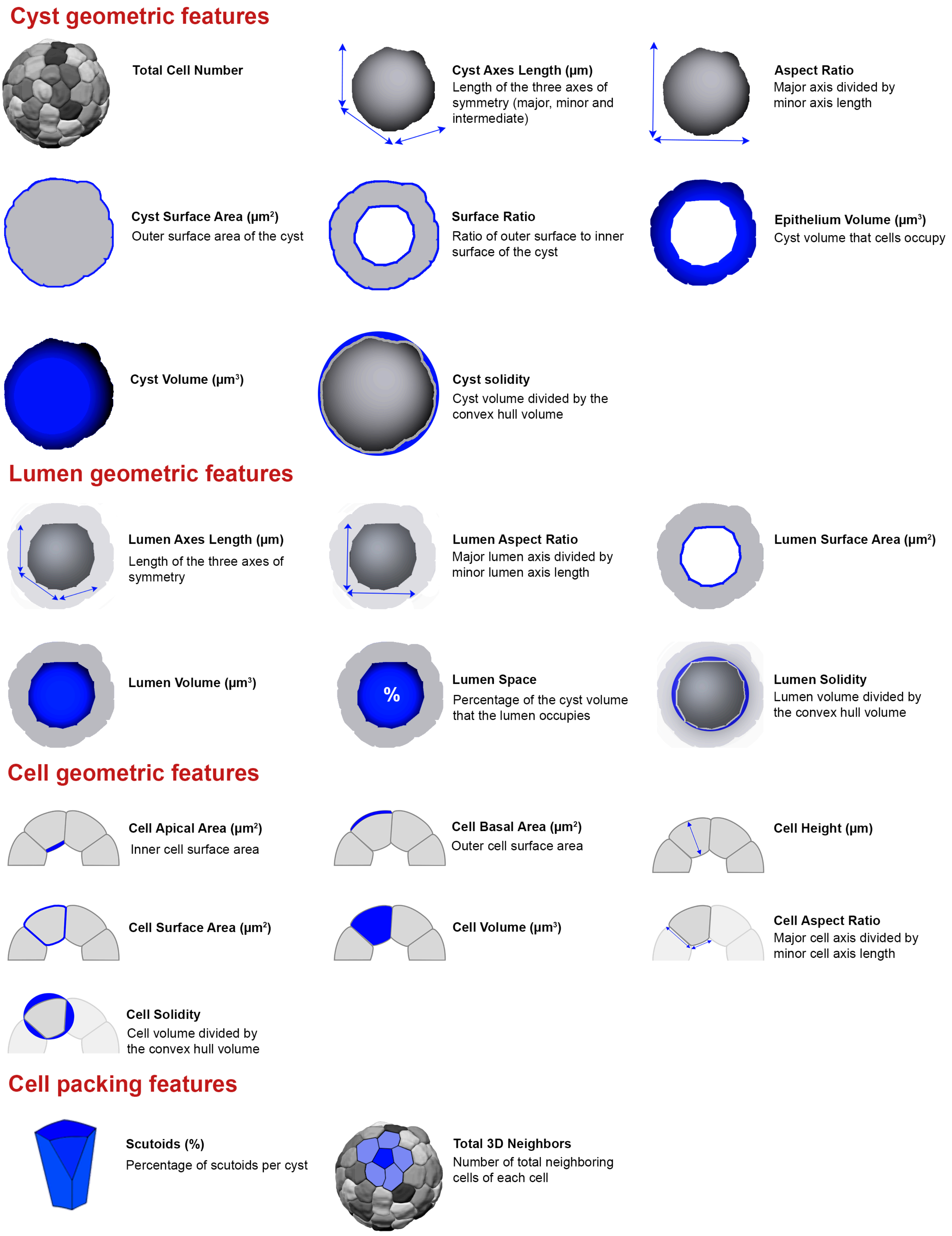

Cyst geometric features
Total Cell Number
Cyst Axes Length (µm)
Length of the three axes of symmetry (major, minor and intermediate)
Aspect Ratio
Major axis divided by minor axis length
Cyst Surface Area (µm²)
Outer surface area of the cyst
Surface Ratio
Ratio of outer surface to inner surface of the cyst
Epithelium Volume (µm³)
Cyst volume that cells occupy
Cyst Volume (µm³)
Cyst solidity
Cyst volume divided by the convex hull volume
Lumen geometric features
Lumen Axes Length (µm)
Length of the three axes of symmetry
Lumen Aspect Ratio
Major lumen axis divided by minor lumen axis length
Lumen Surface Area (µm²)
Lumen Volume (µm³)
%
Lumen Space
Percentage of the cyst volume that the lumen occupies
Lumen Solidity
Lumen volume divided by the convex hull volume
Cell geometric features
Cell Apical Area (µm²)
Inner cell surface area
Cell Basal Area (µm²)
Outer cell surface area
Cell Height (µm)
Cell Surface Area (µm²)
Cell Volume (µm³)
Cell Aspect Ratio
Major cell axis divided by minor cell axis length
Cell Solidity
Cell volume divided by the convex hull volume
Cell packing features
Scutoids (%)
Percentage of scutoids per cyst
Total 3D Neighbors
Number of total neighboring cells of each cell

**Figure S4. Qualitative definition of the geometric and packing features describing the cyst architecture, Related to Figure 2.**

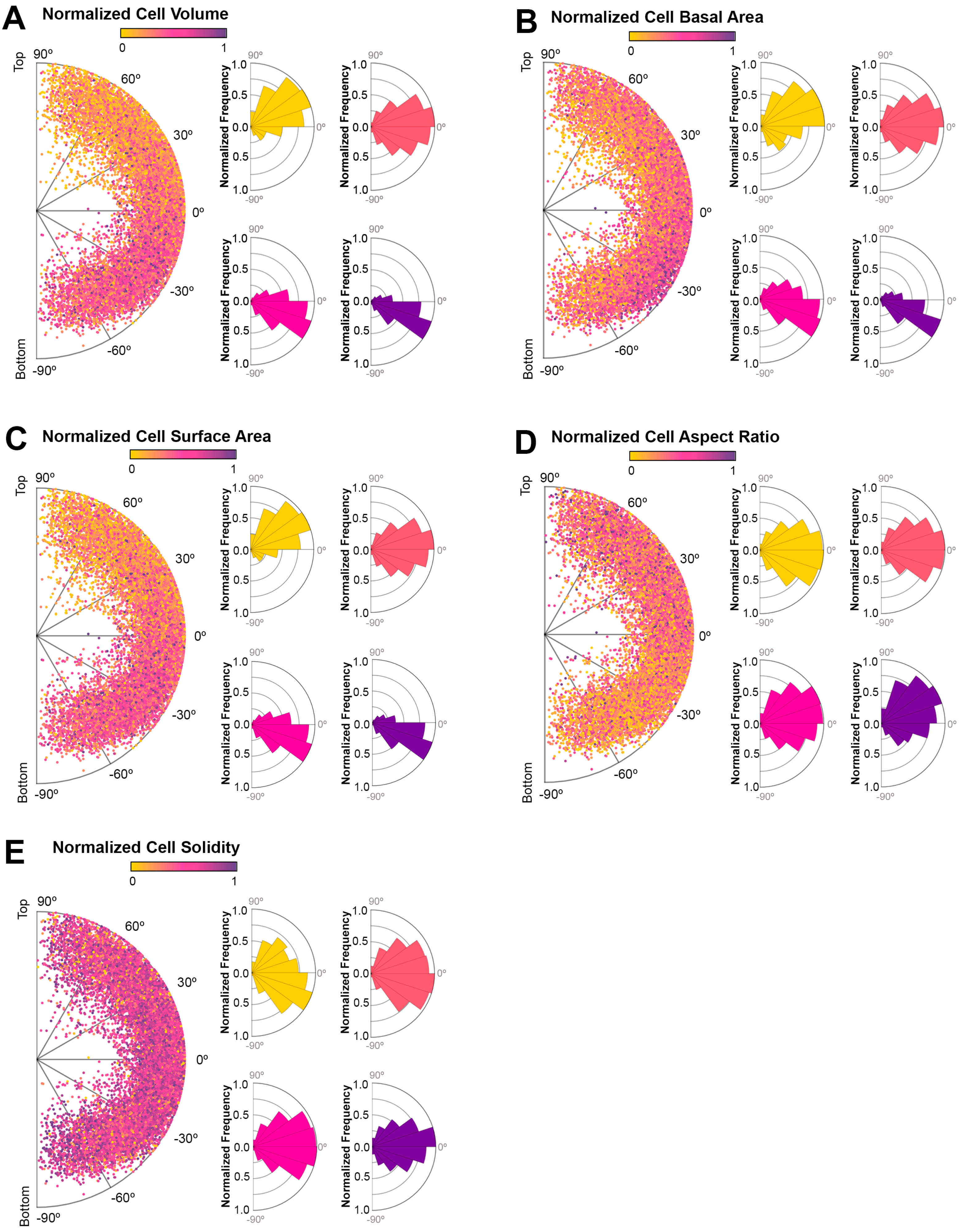
A
Normalized Cell Volume
0
1
Top
Bottom
90º
60º
30º
0º
-30º
-60º
-90º
Normalized Frequency
1.0
0.5
0.0
0.5
1.0
B
Normalized Cell Basal Area
C
Normalized Cell Surface Area
D
Normalized Cell Aspect Ratio
E
Normalized Cell Solidity

**Figure S5. Examples of spatial patterns in the distribution of geometric feature values, Related to Figure 2.**

Same representations as **Fig. 2E**, **2G** to map the features: cell volume (**A**), cell basal area (**B**), cell surface area (**C**), cell aspect ratio (**D**) and cell solidity (**E**).

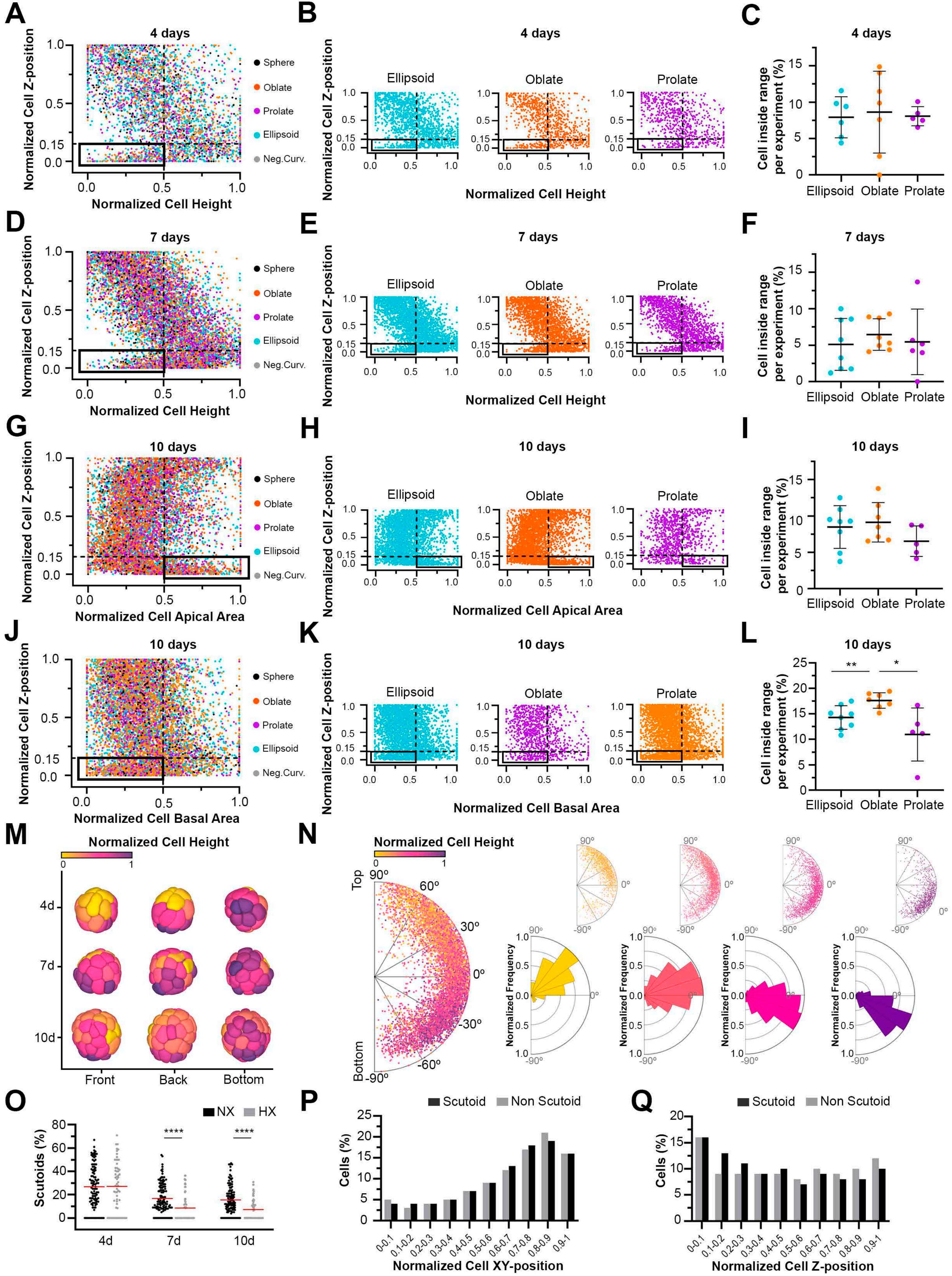

A
4 days
Normalized Cell Z-position
Normalized Cell Height
Sphere
Oblate
Prolate
Ellipsoid
Neg.Curv.
B
4 days
Ellipsoid
Oblate
Prolate
C
4 days
Cell inside range per experiment (%)
D
7 days
E
7 days
F
7 days
G
10 days
Normalized Cell Apical Area
H
10 days
I
10 days
J
10 days
Normalized Cell Basal Area
K
10 days
L
10 days
M
Normalized Cell Height
4d
7d
10d
Front
Back
Bottom
N
Normalized Cell Height
Top
Bottom
Normalized Frequency
O
NX
HX
Scutoids (%)
4d
7d
10d
P
Scutoid
Non Scutoid
Cells (%)
Normalized Cell XY-position
Q
Scutoid
Non Scutoid
Cells (%)
Normalized Cell Z-position

**Figure S6. The cyst morphology and the hypoxia can affect the cartography of cell geometric features, Related to Figure 2 and Figure 3.**

**A-L**) The cyst morphology affects the cartography of cell geometric features. Same representations as **Fig. 2H-2J** to explore the cell Z-position versus cell height at 4-day (**A-C**) and 7-day cysts (**D-F**), cell apical area at 10-day cysts (**G-I**) and cell basal area at 10-day cysts (**J-L**). **M-Q**) Hypoxia can induce cell morphology patterns changes in MDCK cysts. **M**, **N**) Same representations as **Fig. 2D-2G** to map the feature "cell height" in hypoxic (1% $O_2$) MDCK cysts. 7729 cells from 206 segmented cysts were represented. **O**) Scatterplots for quantification of the percentage of scutoids per cyst in normoxic (NX, black) and hypoxic (1% $O_2$) (HX, grey) cysts at 4, 7 and 10 days. The red bar represents the average of the dataset. Data from more than 60 segmented cysts at each time point, from at least 3 independent experiments. "****" indicates $p \leq 0.0001$ (Mann-Whitney U test). **P**, **Q**) Same representations as **Fig. 2L**, **2M** to map the scutoidal (black) and non-scutoidal (grey) cells along XY (**P**) and Z axes (**Q**) in hypoxic (1% $O_2$) cysts. Percentage of all cells from the 206 segmented hypoxic cysts.
